# Phylogenomic framework and virulence gene boundaries of emerging Shiga toxin-producing *Escherichia coli* O118 informed by the comprehensive profiling of 359 O118 genomes

**DOI:** 10.1101/2025.04.29.651274

**Authors:** Irvin Rivera, Sara S.K. Konig, Armando L. Rodriguez, Joseph M. Bosilevac, Mark Eppinger

**Author notes:** Corresponding author: Mark Eppinger, PhD, MS, Department of Molecular Microbiology and Immunology (MMI) & South Texas Center for Emerging Infectious Diseases (STCEID), University of Texas at San Antonio (UTSA), One UTSA Circle, San Antonio, TX, 78249-0600.

## Abstract

Non-O157 Shiga toxin-producing *Escherichia coli* (STEC), particularly the O118 serogroup, are emerging pathogens linked to severe foodborne illnesses, including hemolytic uremic syndrome. The hallmark of STEC virulence is the production of a potent phage-borne cytotoxin, often accompanied by the locus of enterocyte effacement (LEE). This study explores the genomic landscape, virulence factors, and resistance traits of O118 STEC. We analyzed 357 publicly available O118 genomes across ten H-antigens and included two clinically significant O118:H16 STEC strains sequenced to closure. Pangenome assessment and core genome multilocus sequence typing (MLST) based on 4,160 shared genes revealed phylogenetic clustering by H-type and delineated distinct STEC-phylogroups, alongside relationships to non-STEC pathovars such as uropathogenic *E. coli* (UPEC), enteropathogenic *E. coli* (EPEC), and enterotoxigenic *E. coli* (ETEC). Identified STEC phylogroups encompassed H6, H12, H16, and H2 strains with diverse Shiga toxin (*stx*) profiles (*stx_1a_, stx_2a_, stx_2b_, stx_2c_, stx_2f_*). A subset of H2-STEC lacked *stx*, suggesting potential secondary phage loss. Most STEC groups harbored the locus of enterocyte effacement (LEE). Further, a strong correlation was observed between H-antigens and *eae* subtypes, with specific pairings such as H6/*eae-ι*, H16/*eae-β*, and H2/*eae-ε*. Horizontally acquired pathogenicity islands—including O-island 122 in H16 strains and a novel pathogenicity-associated island carrying antibiotic resistance—along with other loci related to colonization and interbacterial competition, further enhance these strains’ virulence potential. Our findings underscore the genetic diversity and virulence potential of O118 STEC. Understanding phylogroup-specific traits and resistance markers is crucial for effective surveillance and public health interventions.

**Importance:** Shiga toxin-producing *Escherichia coli* (STEC) are a major public health concern, responsible for illnesses ranging from diarrhea to life-threatening kidney damage. Among non-O157 STEC, serogroup O118 is increasingly recognized as an emerging STEC lineage. This study provides the most comprehensive genomic analysis to date of O118 STEC, offering critical insights into their pathogenome, virulence traits, and antimicrobial resistance inventories. Notably, we identified a novel 74 kb pathogenicity island in clinical strain H16-12089 that encodes a combination of multidrug resistance genes, virulence factors, and interbacterial competition systems. Understanding the genetic makeup and virulence potential of these pathogens is essential for improving surveillance, risk assessment, and public health interventions.

## 1. Introduction

*Escherichia coli* O118 is an emerging foodborne Shiga toxin-producing *E. coli* (STEC) serotype with a bovine zoonotic reservoir (1). Infections can cause severe human gastrointestinal disease and may progress to life-threatening complications, such as hemorrhagic colitis and hemolytic uremic syndrome (HUS) (2, 3). Vulnerable populations, such as children and the elderly, are particularly affected (4–6).

This underscores the importance of monitoring this emerging serotype in both veterinary and public health contexts and assessing its virulence potential and phylogenetic boundaries (4, 7). *E. coli* are historically classified by their somatic O– and flagellar H-antigens (8–10). Serotype O157:H7 is the dominant causative agent of STEC disease in the U.S. (11–15). However, the incidence of non-O157 infections has steadily increased (16–19). Emerging serogroups O26, O45, O103, O111, O121, and O145 account for most clinical non-O157 STEC infections in the US and are colloquially referred to as the “Big Six” (16, 20, 21). These accounted for 83% of non-O157 STEC infections in the United States from 2000 to 2010 (22–25). As diagnostic testing increased, non-O157 STEC, namely the “Big Six”, surpassed the previously dominant O157:H7 lineage in 2014 (21, 26). Between 2021 and 2023, non-O157 infections accounted for 168k annual cases, compared to 96k O157 cases (22, 27).

The first reported human infection with *stx*_1_-positive O118:H2 STEC occurred in 1996 during a school outbreak in Japan (28) linked to salad (29). Since then, strains of O118 (including its related O151 subgroup) (30, 31) have been recognized as an emerging and human pathogenic STEC lineage, many of which possess Shiga toxins, intimin (*eae*), and hemolysins (29, 32, 33). The carried ΦStx prophages feature different combinations of *stx*-suballeles (12, 34) that can also form hybrid toxins (35). Both *stx*_1_ and *stx*_2_ are potent translation inhibitors (29, 34, 36–41) and their alleles have been documented in O118 (6, 32, 42). Stx_2_, particularly *stx_2a,_* and *stx_2d,_* have been associated with elevated cytotoxicity (43–46) compared to the morbidity caused by *stx_1a_*_+_ STEC (8, 17, 47, 48). However, Stx_1_ has been shown to exhibit increased cytotoxicity in Vero cells compared to Stx_2a_ (44, 49, 50). Adding further complexity, Stx_1_ has been reported to reduce the overall toxicity of Stx_2a_, potentially through competitive binding or interference at the receptor level (44). Besides the pathovar-defining Shiga toxin, another notable virulence factor include the locus of enterocyte effacement (LEE). This pathogenicity island encodes a type III secretion system (T3SS) and effectors, the outer membrane adhesin intimin (*eae*), and its translocated receptor (*tir*) (9, 29, 51, 52). The latter is also carried by enteropathogenic *E. coli* (EPEC) (53). Antimicrobial (AR) and multidrug-resistant (MDR) isolates are described for O118 STEC (29, 32, 54, 55). STEC also carry serogroup-specific virulence plasmids and an array of other plasmids with diverse functions (56–58). Serogroup O118 also contains EPEC, encoding intimin adhesins forming characteristic attaching and effacing lesions (4, 28); Enteroaggregative *E. coli* (EAEC) which form biofilm-like aggregates (4, 16, 59–61); and extraintestinal pathogenic *E. coli* (ExPEC) that are capable of causing severe infections outside the gastrointestinal tract, such as urinary tract infections (UTIs) and bloodstream infections (28, 62). In this study, we provide a comprehensive framework for O118 *E. coli* with a focus on emerging STEC O118. The gathered information on virulence gene content and genome makeup provide a foundation to delineate evolutionary and pathovar boundaries of these emerging and human pathogenic non-O157 lineages.

## 2. Materials and Methods

### Bacterial strains analyzed in this study

For this study, we analyzed 359 O118 genomes, including three closed genomes, 357 publicly available genomes retrieved from the NCBI Pathogen Detection Isolate Browser (https://www.ncbi.nlm.nih.gov/pathogens/isolates/) (July 2024), and two clinical O118 genomes from our STEC collection that were sequenced to closure. Strain-associated metadata and genome statistics can be found in **Table S1**.

### Genome sequencing, assembly, and annotation

Strains were cultured overnight at 37°C with shaking at 220 rpm in lysogeny broth (LB) (Thermo Fisher Scientific, Asheville, NC, USA). To maximize total genomic DNA (gDNA) yields, bacterial overnight cultures were diluted to OD_600_ of 0.03 in fresh LB medium and grown at 37°C with shaking at 220 rpm to mid-log phase (OD_600_∼0.5). Total gDNA was extracted using the Monarch HMW DNA Extraction Kit (New England Biolabs, Ipswich, MA, USA). Genomes were sequenced to closure using Nanopore long-read technology (Oxford Nanopore, UK). Sequencing libraries were prepared with Rapid Barcoding Kit (RBK-114) according to the manufacturer’s instructions and sequenced on the PromethION platform on R10.4.1.Flow Cell (FLO-MIN114). Reads in the fastq format were imported into Galaxy v.22.05 (63).

Default parameters were used for all software unless specified otherwise. Fastq reads were QC-ed using FastQC (v.0.74+Galaxy0) (http://www.bioinformatics.babraham.ac.uk/projects/fastqc) and assembled with Flye (v.2.9.3) (64). The resulting contigs were evaluated with QUAST (v.5.2.0 + Galaxy1). The chromosomal *dnaA* and plasmid *repA* genes were designated as the zero point of the closed molecules prior to annotation using the NCBI Prokaryotic Genome Annotation Pipeline (PGAP) (65). Genome assemblies were profiled with SeqKit v2.8.2 to obtain genome statistics (66, 67).

### Phylogenomic analyses and MLST-schemas

The O118 genomes, along with the sequence of K-12 substrain MG1655 (68, 69), were imported into Ridom SeqSphere+ (v.8.3) (Ridom GmbH, Münster, Germany) for core genome (cg) and targeted Multilocus Sequence Typing (MLST) (70–72). The Sequence Type (ST) was determined according to the EnteroBase schema (73). Allele sequences for the Achtman scheme, targeting seven housekeeping genes (*adk*, *fumC*, *gyrB*, *icd*, *mdh*, *purA*, and *recA*), were accessed on the EnteroBase website (https://enterobase.warwick.ac.uk/species/ecoli/download_7_gene). A cgMLST schema was developed using the closed chromosome of K-12 substrain MG1655 (69) as seed as previously described (74). Core and accessory MLST targets were identified according to the inclusion/exclusion criteria of the SeqSphere+ Target Definer. The allele information from the targeted seven-gene schema and the defined core genome gene of the panel strains were used to establish phylogenetic hypotheses using the minimum-spanning method with default settings (75, 76).

### Pathogenome makeup and visualization

Chromosomes were compared with progressive Mauve (77) and visualized in pyGenomeViz-pgv-mauve (78) (https://github.com/moshi4/pyGenomeViz) using seaborne (79). Chromosomes and carried plasmids were compared and visualized in Blast Ring Image Generator BRIG (v.0.95) (80). Serotypes were determined *in silico* using *E. coli* Typer (ECTyper) (https://github.com/phac-nml/ecoli_serotyping) (Galaxy 1.0.0) in Galaxy v.22.05. If the *wzy* gene was reported absent by ECTyper, BLASTn was used to reevaluate its presence and fragmentation status. The ECTyper database was downloaded as a JSON file from GitHub (https://github.com/phac-nml/ecoli_serotyping) and converted into a blast database. The Average Nucleotide Identity (ANI) between all-vs-all 359 genomes was computed with skani (81), the distance matrix clustered by the UPGMA (Unweighted Pair Group Method with Arithmetic Mean) method and visualized using ANIclustermap (78) (https://github.com/moshi4/ANIclustermap). Virulence and antibiotic resistance genes (ARGs) were cataloged with Virulence Factors of Pathogenic Bacteria (VFDB) (82) and ResFinder (https://cge.cbs.dtu.dk/services/ResFinder/) (83), respectively. Pathovars were inferred *in silico* through BLASTp query (84) against a curated VFDB database of *E. coli* pathovar-defining virulence genes (82, 85–92). Ribosomal RNAs and CRISPR/Cas systems were detected with Basic Rapid Ribosomal RNA Predictor (Barrnap) (93) (v.0.7) and CRISPRCasFinder (94) (v4.2.20) in Proksee (95) (v.1.1.0). Anti-phage systems were detected using DefenseFinder (96) (Galaxy v.1.3.0+galaxy0). Prophages were compared and visualized in Easyfig (v.2.2.2) (97). Boundaries and locations of intact, partial, or remnant prophages were identified using PHASTEST (98), followed by manual curation of the ΦStx-prophage and core genome borders by Mauve alignment of the genome to *E. coli* K-12 substrain MG1655 (GenBank accession U00096) (68) in Geneious Prime (v.2024.0.5). Direct repeats caused by phage integration were identified by BLASTn self-alignment and the Repeat Finder plugin (v1.0.1) in Geneious Prime. Replicase subtyping was performed to characterize the ΦStx_1a_-encoding prophages, following the typing scheme introduced by (99). Replicases were typed with EHEC phage replication unit (*eru*) schema (100, 101). The subtypes of phage-borne *stx* were determined by BLASTn against a curated *stx*-suballele database, as previously described (58, 102–104). A hierarchical classification approach was implemented to categorize phage-associated genes from the PHASTEST JSON output. All present phage-associated genes were further curated processing type and name descriptors, along with reported Gene Ontology (GO), for more in-depth functional classification. Genomic islands (GI) were detected with IslandViewer4 (105–107). LEE islands and boundaries were curated, guided by LEE1-LEE4 operon genes *espG and espF,* respectively, and visualized in EasyFig (v.2.2.2) (97). Intimin subtypes were determined by BLASTn against a curated database of published *eae*-subtype sequences, as reported previously (58). Transposable elements and Insertion Sequence (IS) elements, along with inverted repeats, were cataloged through BLASTn (108) against the Transposon Central (TnCentral) database (109), comprised of TnCentral+Integrall+ISFinder entries (110, 111) and ISEScan (v.1.7.2.3+Galaxy0) (112).

### Plasmid makeup and visualization

Plasmids were visualized and decorated in Blast Ring Image Generator BRIG (v.0.95) (80). Plasmid mobility and incompatibility groups were recorded with Mobilome Typer (MOB-Typer) (v.3.0.3+Galaxy0) (113) and ABRicate (Galaxy v.1.0.1; (https://github.com/tseemann/ABRicate) with options ––minid 95 ––mincov 90 using the PlasmidFinder database (108, 114). Using a mash search strategy, phylogenetically related plasmids were identified with the Plasmid DataBase (PLSDB) tool (v.2024.05.31.v2) (115) (116). Serotypes of the plasmid-associated chromosomes molecule were determined *in silico* using ECTyper (https://github.com/phac-nml/ecoli_serotyping) (Galaxy v1.0.0) in Galaxy v.22.05. Virulence genes (ARGs) were cataloged with VFDB (82). Bacteriocins, ribosomally synthesized and post-translationally modified peptides (RiPPs), were recorded with BAGEL4 (http://bagel4.molgenrug.nl/) (117). Genomic islands were detected with IslandViewer4 (105–107).

Mobile Genetic Elements (MGEs) were identified with the mobile orthologous groups database (mobileOG-db) tool (118) in Proksee (95). Transposable elements and IS elements were identified with ISEScan (v.1.7.2.3+Galaxy0) (112) and through BLASTn query (108) against the TnCentral database, comprised of TnCentral+Integrall+ISFinder entries (109–111).

### Pan-Genome computation and visualization

The global gene reservoir of serogroup O118 was computed by re-annotating all genomes with Prokka (v.1.14.6+galaxy1) (119). The generated gff3 files were then subjected to pan-genome analysis using Roary (v.3.13.0+galaxy2) (120). Classification of full-length genes into core, accessory, and shell genes provided information on shared and unique genes according to Roary’s inclusion/exclusion criteria (121), and visualized Phandango (v.1.2.0) (122).

## Results and Discussion

### Genomes of closed STEC O118 genomes

At the time of data collection, three closed serogroup O118 genomes were publicly available, all of which are STEC: H6-strains 2013C-4538 and 2014C-3050 from human stool (123) and H2-strain EC20017429 isolated in 2017 from a gastroenteritis patient (124). To support comprehensive analyses of O118 STEC, we sequenced two clinical H16-isolates, strains 12089 and 12867, to closure using long-read sequencing (20, 125, 126). This approach yielded high-quality closed chromosomes of 5,804,173 and 5,685,925 bp and recovered carried plasmids (**Fig. 1**, **Fig. 2**). Genome statistics and strain-associated metadata are provided in **Table S1**. The average nucleotide identity processing closed and draft O118 genomes was 99.34%, indicative of the highly conserved chromosomal *E. coli* backbone (127, 128) (**Fig. S1**). The STEC strains contained about 1 Mb of additional sequence information compared to nonpathogenic *E. coli* strain K-12 substrain MG1655 (68), included in this comparison. Mobile genome elements are major drivers of pathogenome diversification (38, 129–133). The identified prophages, GIs, and IS elements contributed significantly (24.11 to 28.51%) to the genome content of the closed O118 STEC, comparable to observations in other STEC-lineages (133, 134) (**Table S2**). These included pathovar-defining ΦStx prophages (29, 34, 36–41) and other virulence hallmarks, such as the LEE Pathogenicity Island (29, 52, 135). IS elements have been utilized to delineate distinct phylogenetic lineages in STEC (20, 136–138).

**FIG 1.**
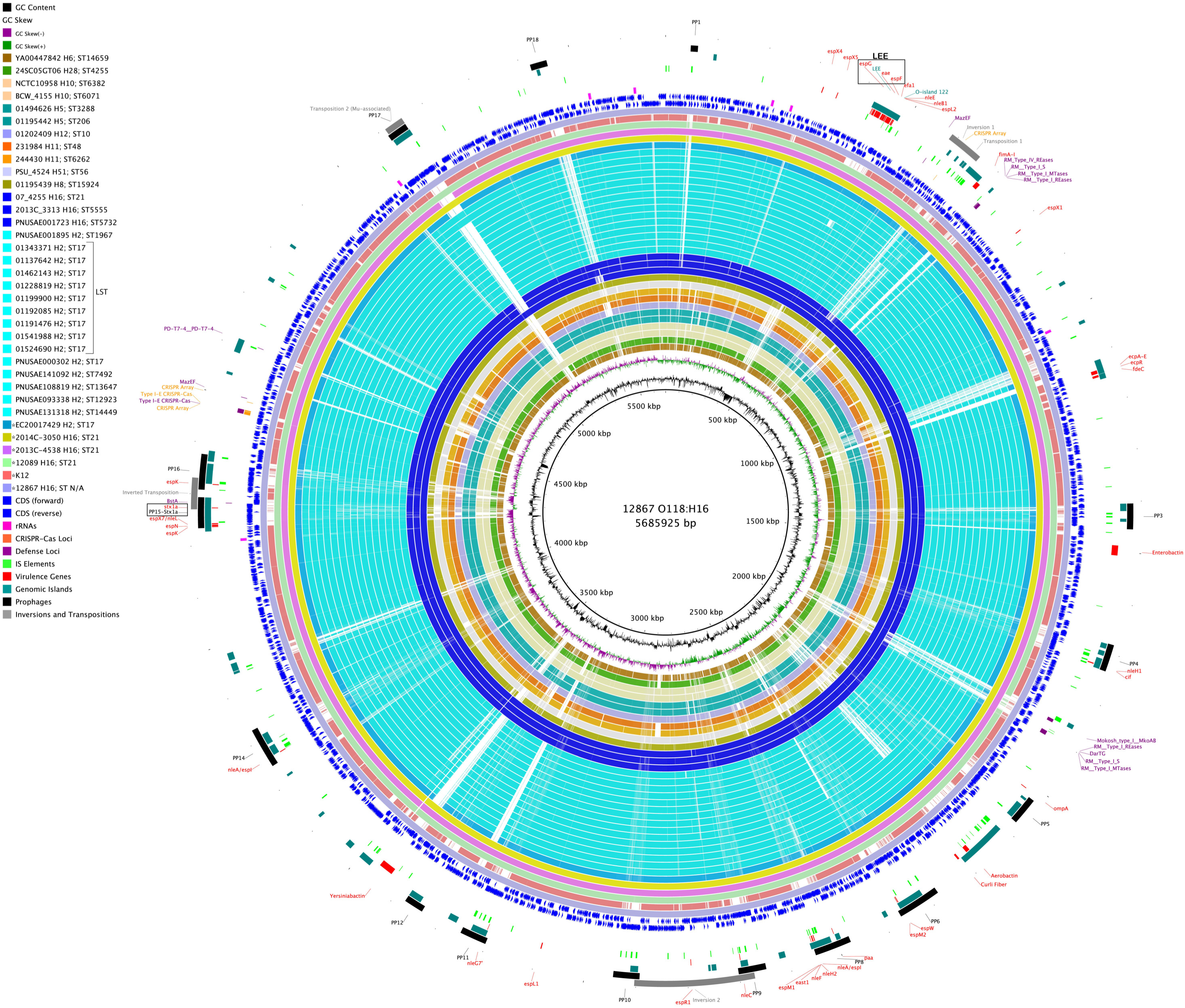
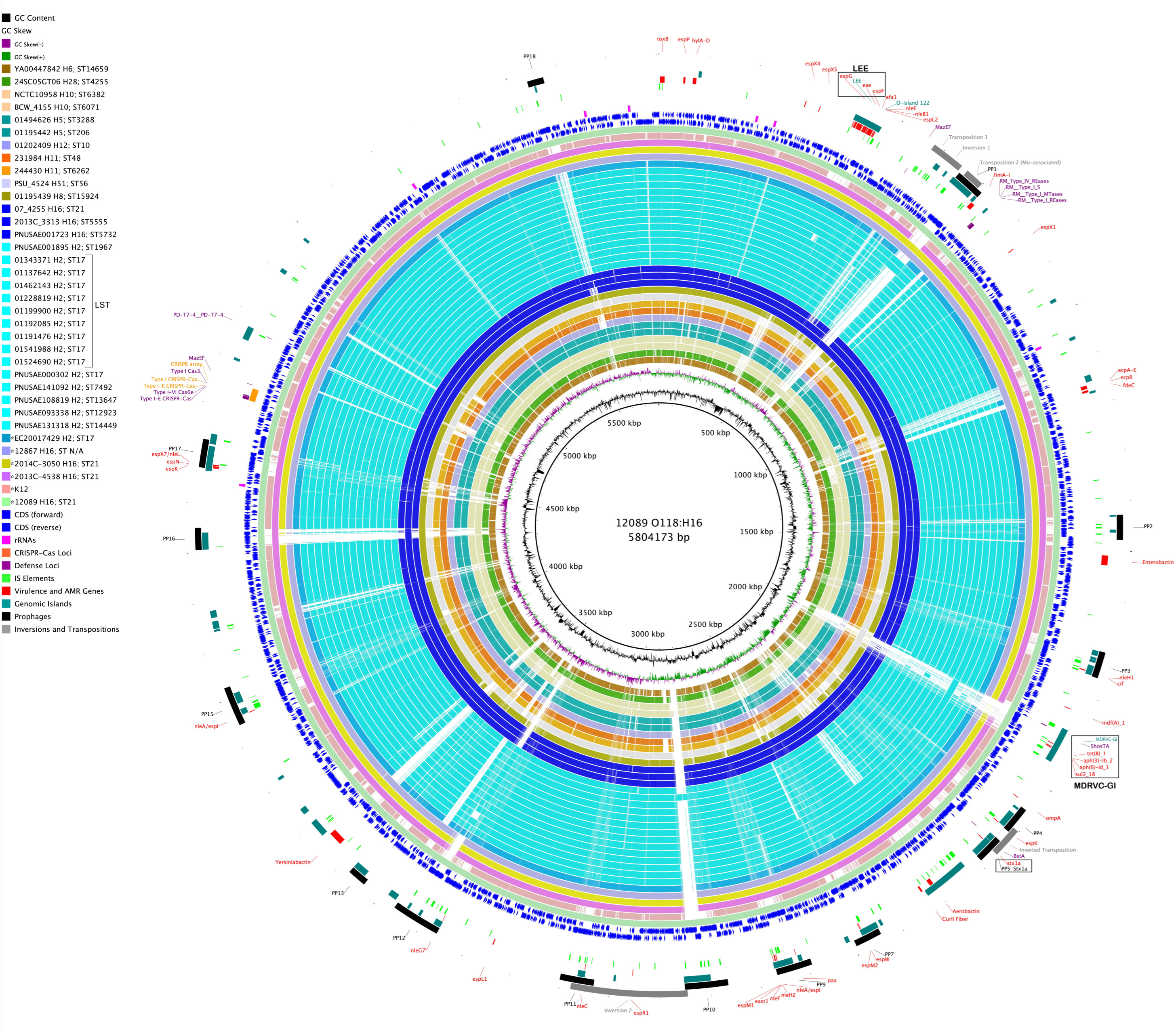
Comparison of O118 STEC chromosomes. BRIG comparison of representative genomes for all identified STEC-phylogroup, considering H-antigen and sequence type, referenced to O118:H16 strains **A)** 12867 and **B)** 12089. *E. coli* K-12 is also included to highlight STEC-specific genome content. Query genomes are color-coded, and their order reflects the inferred phylogenomic relationships. Coding sequences are shown as arrows on the + and – strands, and functional annotations for virulence genes and other loci of interest are highlighted.

**FIG 2.**
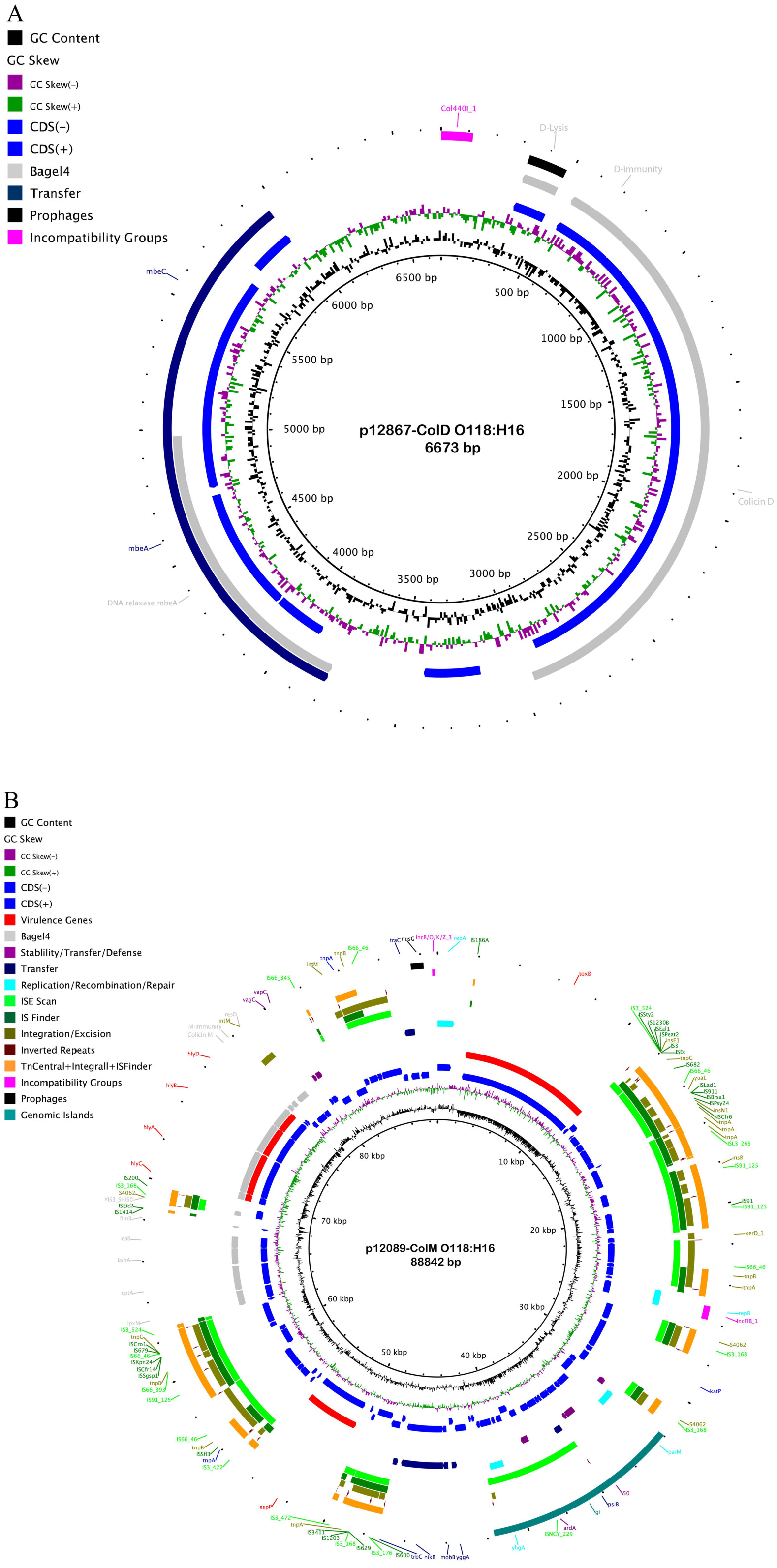
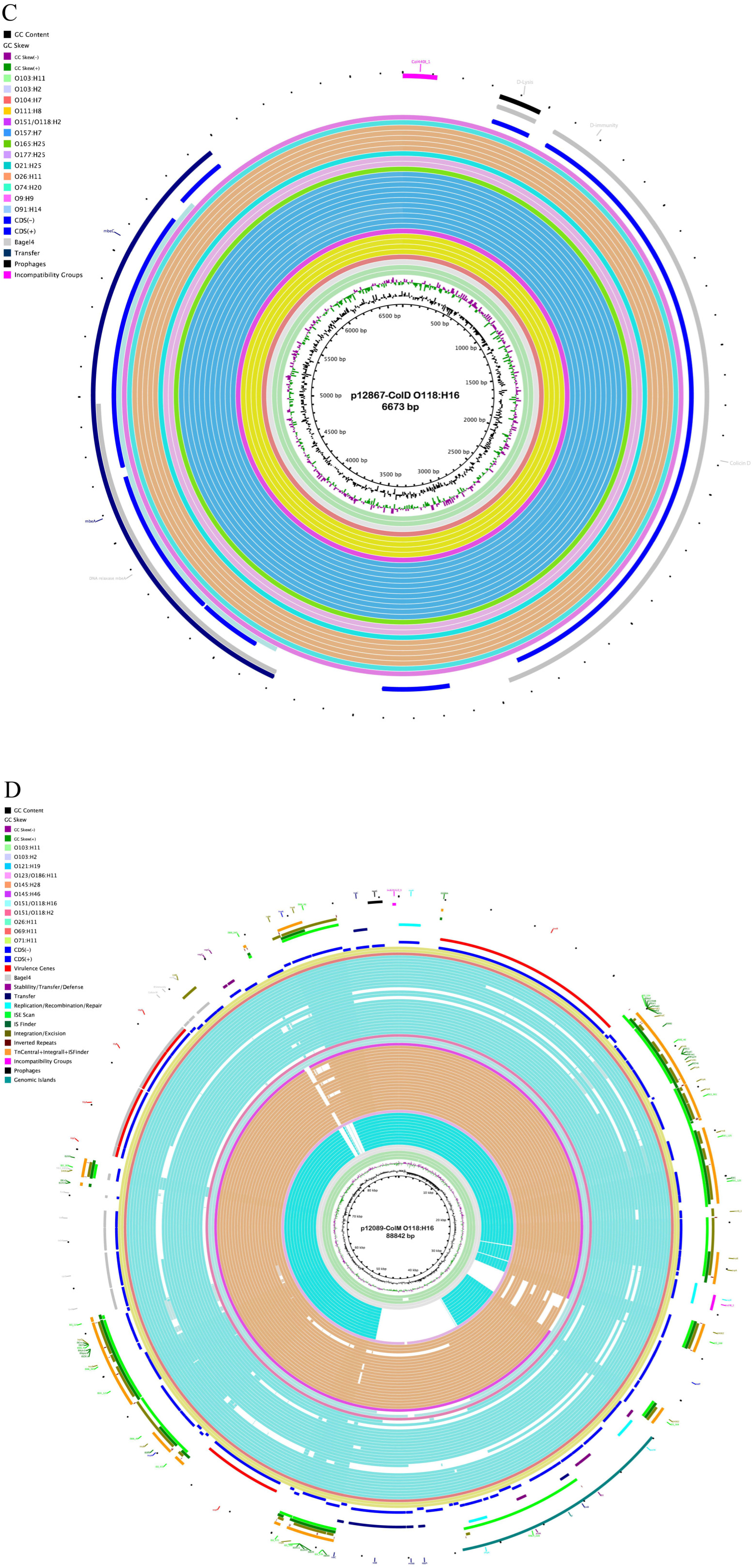
Plasmid inventory of the two sequenced O118 STEC strains. **A)** The small 6,673 bp ColD-colicin plasmid in H2-strain 12867, **B)** The 88,842 bp multifunctional ColM-colicin and virulence plasmid of H16-strain 12809 features an array of virulence and conjugation loci, **C)**, and **D)** Phylogenetically related plasmids were detected in diverse STEC serogroups.

We recorded IS element prevalence and types in the five closed genomes (**Fig. S2**), showing both shared (IS3_168) but also H-antigen-specific IS elements (e.g., IS3_255, IS4_463). Mauve comparison of closed O118 chromosomes revealed largely genome-wide chromosome synteny but identified several inversions, inversion/transposition, and transposition events associated with adjacent or flanking mobility or ribosomal RNA loci (**Fig. S3**) (139–142). The sequenced H16-strain 12867 and H16-strain 12089 are both colicinogenic (**Fig. 2**). Colicins are produced and toxic to *E. coli,* and support the ability of *E. coli* to compete for a shared niche (143). Plasmid p12867-ColD (143), showed the typical organization into D-lysis, D-immunity, and Colicin D (144), along with a mobilizable DNA-relaxase *mbeA* (145). Additionally, the plasmid-borne SOS inhibition gene *psiB* was detected (146). A MASH-based survey identified related plasmids in STEC O157:H7, the Big Six serotypes, and other STEC-associated serotypes (**Table S2-7**). The larger 88 kb plasmid p12089-ColM encodes Colicin M and its associated M-immunity protein (147). However, its other gene content identified it as a virulence plasmid. Notable were hemolysin (*hlyCABD)* (148), serine protease (*espP)* (149), fimbrial regulator *fimB,* and the STEC adherence factor *toxB* (150). Additionally, modulators of lipopolysaccharide and exopolysaccharide biosynthesis (*lpxM*, *icaB*) (151, 152) along with SOS-inhibition gene *psiB* (146), were also present. Approximately 43% of the plasmid sequence are mobility and conjugation loci (153, 154). A MASH-based plasmid survey identified related plasmids among H16 and H2, Big Six and other STEC serogroups (155–157) (**Table S2-7**).

### Comparison of ΦStx prophages in closed O118 STEC

Comparative genomics provided insights into the shared and specific virulence traits in the identified STEC-phylogroups. Bacteriophages target conserved chromosomal loci and undergo evolutionary acquisition, loss, microevolution, collective shaping, gene content, and pathogenicity traits (132, 158, 159). A comparison of the complete ΦStx_1_ prophages extracted from the closed genomes identified two insertion sites: *torS* and an intergenic insertion upstream of *dinI* with the phage boundaries marked by flanking direct repeats (**Fig. 3**, **Table S2-2**). The *torSTR* locus was one of four identified ΦStx_1a_ integration sites in O26:H11 ST21 (160), an intergenic attachment site common in *E. coli* (161). The ΦStx_1a_ prophages show an overall high-degree of sequence identity and syntenic organization. The *torS* ΦStx_1a_ phage of the H2-strain EC20017429 was distinguished by the presence of an IS3-like element and cluster of three hypothetical genes and a kinase, absent in the other H16 strains. The larger ΦStx_1a_ of H16-strain 12867 carried a DUF550 protein, type III effectors EspK and EspN, and a hypothetical protein.

**FIG 3.**
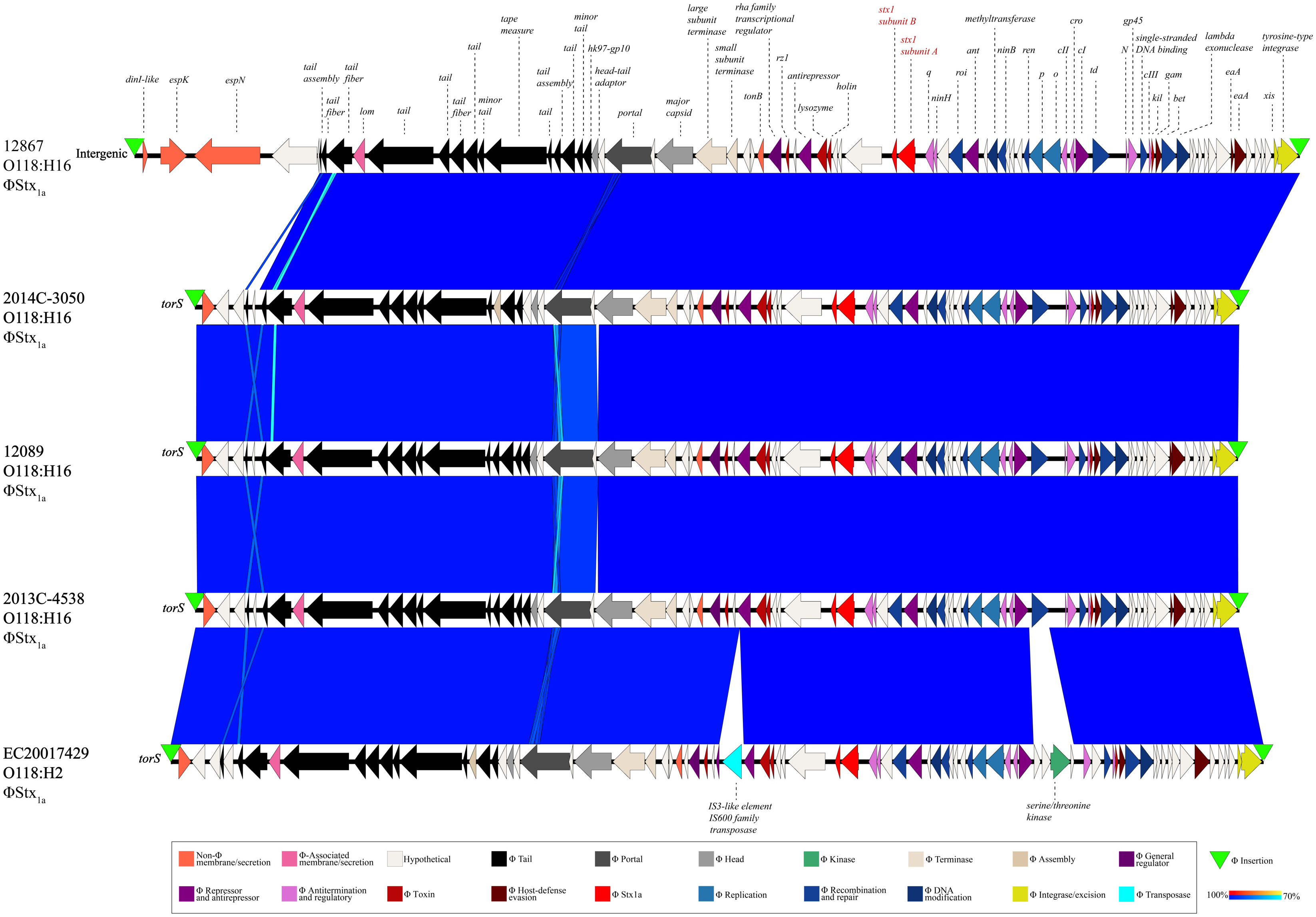
Comparison of ΦStx-prophages in closed O118 STEC. BLASTn-based comparison of the ΦStx_1a_ prophage architectures and gene content. The comparison shows that ΦStx1a phages show considerable genomic plasticity and are inserted at *torS* or intergenically close to the *dinI* locus.

### Locus of Enterocyte Effacement Pathogenicity Island in closed O118 STEC

The Locus of Enterocyte Effacement (LEE) pathogenicity island is carried by diverse STEC serogroups, forming characteristic attaching and effacing (A/E) lesions (52, 162–166). The comparison of LEE islands extracted from the closed O118 STEC genomes (**Fig. 4**) showed they were highly conserved and organized in five polycistronic operons (LEE1 to 5). These encoded T3SS regulators and effectors (167).

**FIG 4.**
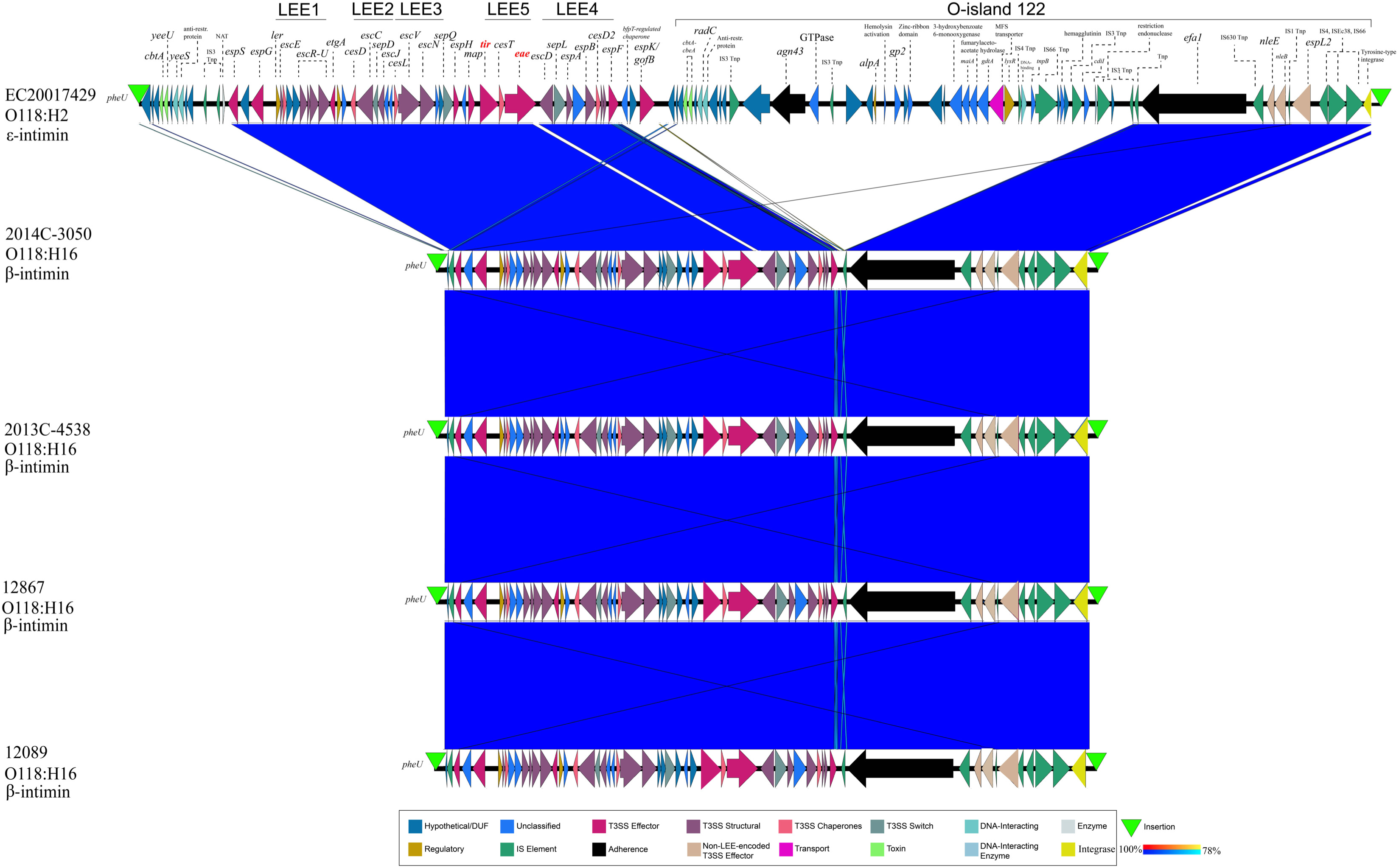
LEE islands of closed O118 STEC. BLASTn-based comparison of LEE islands inserted into the *pheU*. The organization of the LEE1 to LEE5 operons and O-island 122, disrupting the LEE island in H2-strain EC20017429, is indicated. The order of strains reflects their inferred phylogenomic position.

All were integrated at *pheU* tRNA, a known LEE integration site in *E. coli* (168–172). The H2 strain was distinguished by an adjacent O-island 122 within the LEE island boundaries marked by 17 bp direct repeats (**Table S2-4**). This island encodes non-LEE-encoded (Nle) effectors and transport and regulatory elements that enhance *E. coli* pathogenicity by promoting intracellular survival, adhesion, immune modulation, and enterotoxin production (173, 174). Its carriage has been associated with highly virulent STEC, causing HUS (174, 175). The H16 phylogroup, represented by strains 2014C-3050, 2013C-4538, 12867, and 12089, features the β-intimin subtype, whereas the ε-intimin subtype is present in the H2 strain EC20017429 (**Fig. 4**).

### Pathogenicity associated multi-drug resistance, virulence, and competition island

In strain H16-12089, a novel 74 kb Resistance, Virulence, and Competition (MDRVC) Island with a complex modular architecture was discovered inserted into tRNA-Ser (153) (**Fig. 5**). This island featured a tyrosine-type integrase (176), with borders marked by direct repeats (177) (**Table S2-4**). A core component of this island was the MDR-element AMR-SSuT, which has been previously described in STEC O157, though the genomic context was unclear (178). This island conferred resistance to tetracycline, streptomycin, and sulfonamide. Embedded within this element was streptomycin resistance transposon Tn5393.2 (179), itself disrupted by an ISVsa5-flanked Tn10 (180), introducing tetracycline (181, 182). The *sul2* locus, however, is located outside this Tn5393.2-Tn10 composite, proximate to IS66-like elements, previously linked to antimicrobial resistance in *E. coli* (183). Prominently featured were genes that aid interbacterial competition, such as a *cdiA*-mediated contact-dependent inhibition (184) associated with VENN motif pre-toxin (185) and the *cbtA-cbeA* toxin-antitoxin system (186). A broad array of virulence loci was catalogued; among these was the hemagglutinin adhesin (184) and the hemolysins secretion gene (*fhaC*), which likely interacts with the hemolysin on p12089-ColM (**Fig. 2**). Further diverse regulator genes were associated with stress-response. This included *lysR* (187), a DEAD/DEAH-box helicase (188), *alpA* (189), and a bacteriophage defense gene *gp2* (190), along with membrane modeling genes, patatin-like phospholipases and dynamin-like protein *rdcB* (191, 192). DNA-modification enzymes support genomic integrity through repair (*radC*) or endonuclease-mediated defense (193, 194). Altogether, the carriage of this multifunctional GI by strain 12809 suggests increased pathogenic potential and fitness, promoting virulence, antimicrobial resistance, stress adaptation, and also genomic stability.

**FIG 5.**
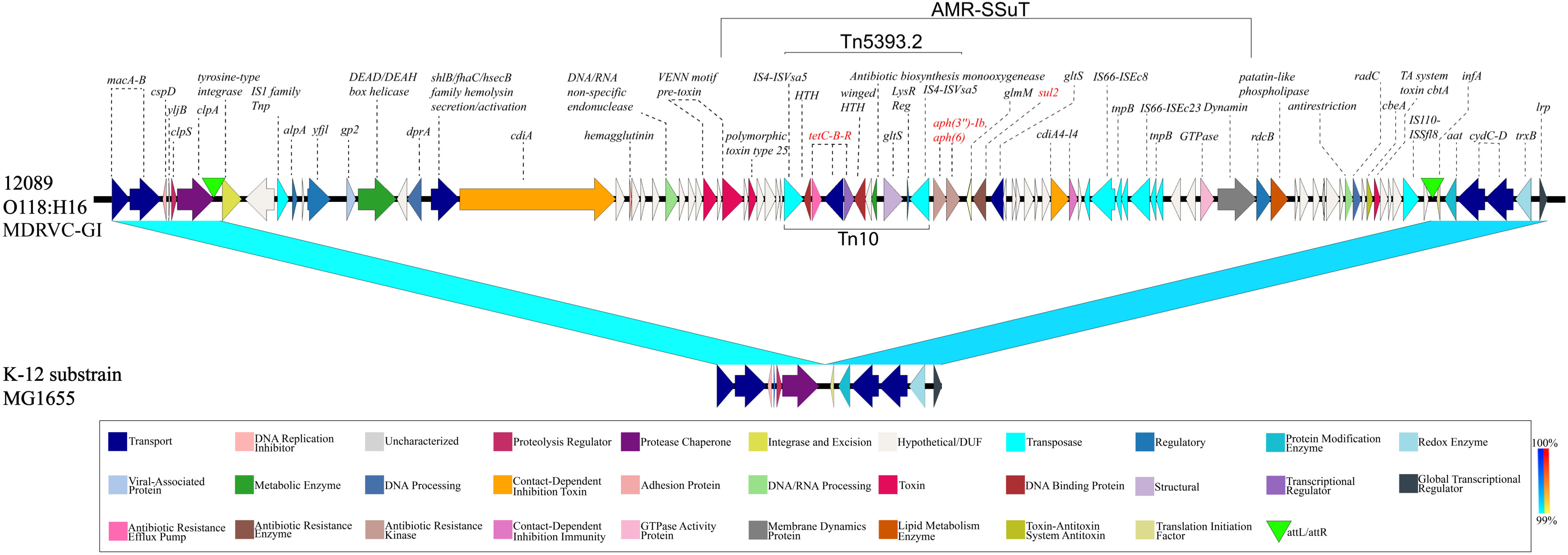
Modular architecture of the MDRVC Island. This newly discovered 74kb multifunctional island, inserted into the *tRNA-Ser* of H16-strain 12089, features a complex modular architecture. It integrates an antimicrobial multi-drug resistance cassette with gene loci, promoting virulence, interbacterial competition, stress resilience, and overall fitness.

### Phylogenomic framework and pathovar boundaries of serogroup O118/O151 STEC

To support broad-scale analysis, we included all publicly available serogroups O118 and O151 genomes retrieved from the NCBI Pathogen Detection Isolate Browser (July 2024), which were all in the draft stage. These were differentiated by subtle genetic and antigenic changes in their O-antigen sequences and linkages (195–197). Among the 359 analyzed strains, 358 possessed the O151-*wzy* variant, while a single strain carried O118-*wzy* (**Table S1-2**). Strain-associated metadata and genome statistics of this expanded strain panel can be found in **Table S1**. This panel of 359 O118 genomes features ten H-antigens, as determined by *in silico* flagellin (*fliC*) subtyping (198, 199). The shared gene inventory was determined at 4,160 genes, comprised of 2,437 core and 1,723 accessory loci, indicative of the genome plasticity found in heterogenous O118 serogroup *E. coli* (**Fig. 7**, **Fig. S4**, **Table S4**). The ANI of 96.32% was computed through an all-vs-all comparison of the 359 O118 genomes (128). The UPGMA-clustering of ANI values recovered the different H-phylogroups (**Fig. S1**). The majority belonged to H2 (n=242), followed by H16 (n=93) and H11 (n=11), with relatively few available representative genomes of H10 and H12 (n=3), H5 and H28 (n=2), and a single strain for H6, H8, and H51 (**Table S1-2**). To investigate the phylogenomic and virulence boundaries of emerging O118 STEC among the heterogenous serogroup O118/O151 strains, we established a phylogenomic framework informed by targeted and core genome MLST (**Fig. 6**, **Fig. 7**). The tree topology partitions the isolates by H-antigen and ST. Pathovars were inferred *in silico* from their VFDB virulence gene profiles. The STEC phylogroups among serogroup O118 were comprised of H2, H6, H12, and H16 strains. Non-STEC pathovars belonged to EPEC (H5, H11), ETEC (H10, H51), UPEC (H28, H10, H12, H11), and a single strain containing virulence genes associated with both ETEC and EPEC (**Fig. 7**, **Table S1-1**, **Table S3**). Notably, H10 antigen belonged to EPEC and UPEC, which formed distinct clusters, also evident in the ANI heatmap (**Fig. S1**).

**FIG 6.**
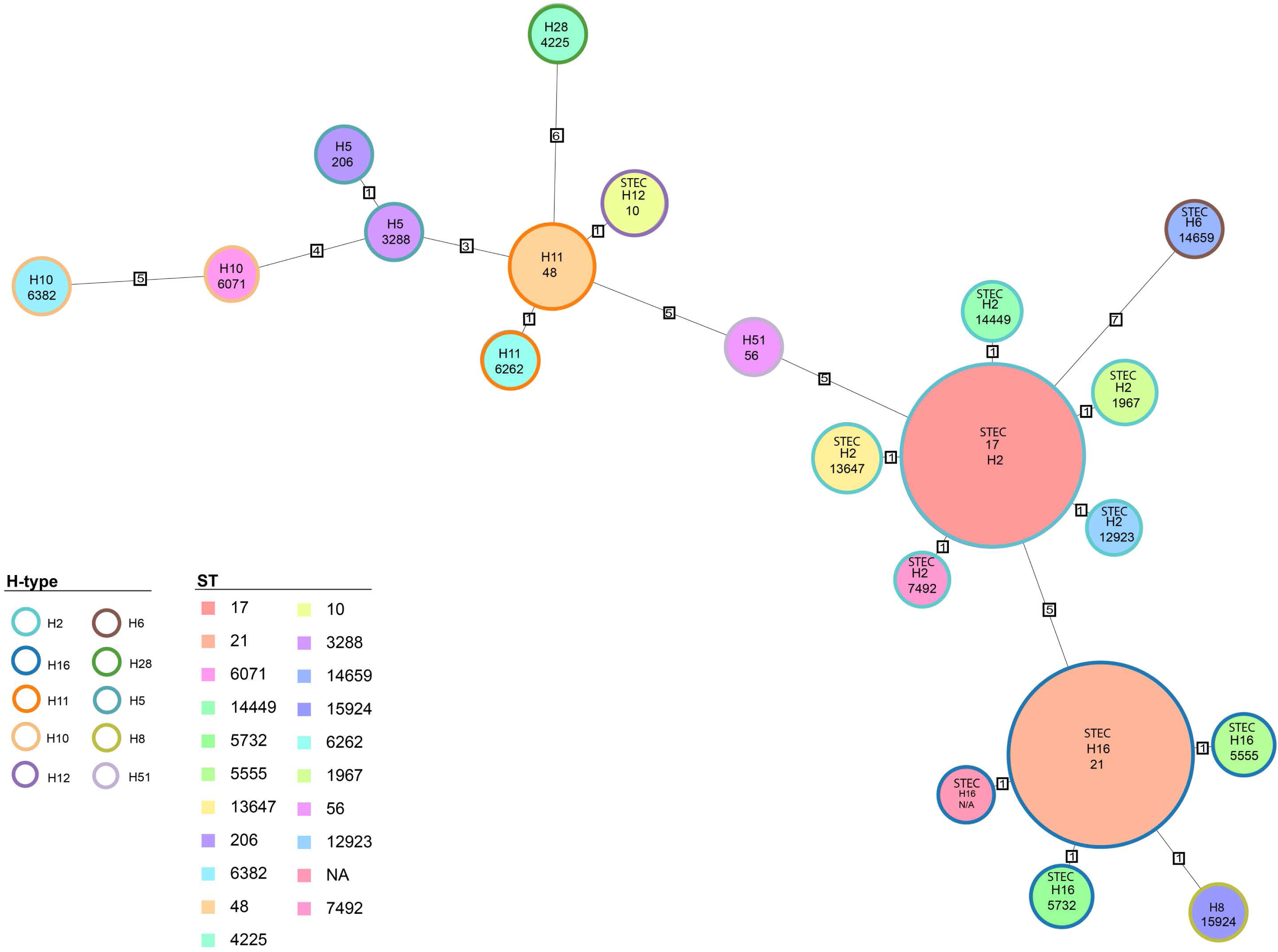
MLST-inferred phylogeny of serogroup O118. The relatedness of O118 strains was determined using MLST in Ridom SeqSphere+ through both targeted seven-gene MLST accessed in the EnteroBase website. Numbers on connecting branches indicate the number of genes with differing allele status. The shared gene inventory of 4,160 genes comprises 2,437 core and 1,723 accessory loci according to the inclusion/exclusion criteria of the SeqSphere+ Target Definer. Circle size corresponds to the number of isolates with identical ST.

**FIG 7.**
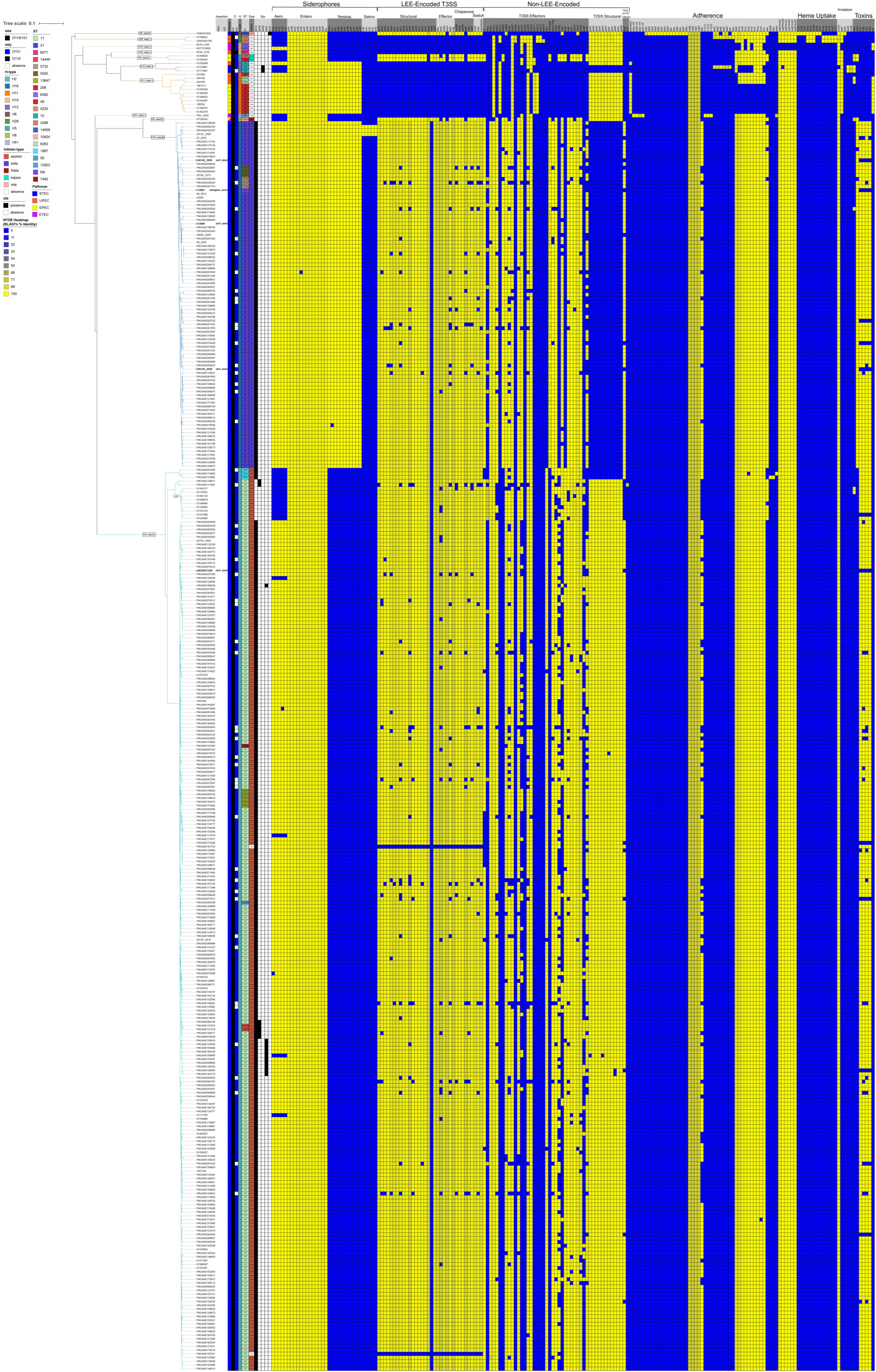
Phylogeny and distribution of virulence genes. The relatedness of analyzed O118/O151 strains was determined using cgMLST in Ridom SeqSphere+ using the closed chromosome of *E. coli* strain K12 substrain MG1655 as seed, complemented by the 357 publicly available strains and two closed genomes strains 12089 and 12867. The shared gene inventory was determined at 4,160 genes. These comprise 2,437 core and 1,723 accessory loci according to the inclusion/exclusion criteria of the SeqSphere+ Target Definer. The topology and clusters indicate a strong association between H-antigen, *eae*-subtype, and ST in LEE+ isolates. A total of 199 distinct virulence genes were cataloged, of which 16 are shared. The STEC-pathogroups are comprised of H-antigens 6, 12, 16, and 2 and carry the *stx*_1a,2a,2b,2c,2f_ suballeles.

### Correlation between Stx status, H-antigens, and intimin subtypes

For STEC, the following *stx/eae*-genotype profiles were recorded: H6 (*stx_2f_*, *eae-ι*), H12 (*stx_2b_*, *eae*(-)), H16 (*stx_1a_*, *eae*-*β*) and H2 (*stx_1a_*, *eae*-*ε*; *stx_2a_*, *eae*-*ε*; *stx_1a_, stx_2c_*, *eae*-*ε*; *stx_1a_, stx_2a_*, *eae*-*ε* and *stx_1a_, stx_2c_*, *eae*-*ε*) (102). The best-represented STEC groups in the analyzed strain panel were H16 (n=93) and H2 (n=242). H16 STEC were genetically uniform (*stx_1a_*, *eae*-*β*) compared to H2 STECs with significant toxicity profile heterogeneity. The dominant Stx-genotype was *stx_1a_+* (93%), followed by isolates with combinations of *stx_2a_* (*stx_1a_+stx_2c_+,* n=11) or *stx2c* (*stx_1a_+stx_2a_*+, n=5), and *stx_2a_+* only (n=2). Within the dominant H2 STEC, we further identified a distinct *stx*(-) sublineage. Further analyses of the H2 ΦStx_1a_ phage insertion site at *torS* (**Fig. 3**) detected phage remnants (**Fig. S5**). We thus speculate that this lineage evolved from ST-17*, stx_1a_+* ancestors and represents Lost Shiga Toxin (LST) isolates (125). Their closest relatives were *stx_2a_+* strains that similarly lack *stx_1a_+*. However, the draft status of these genomes hindered further investigation into Φthe Stx insertion locus at *torS* to evaluate the ancestral or evolved ΦStx-status (**Fig. S5**). We thus speculate that this STEC lineage may have secondarily acquired *stx_2a_*, essentially transitioning between *stx*(+/−) states (125, 200, 201). Similarly, the LEE-H12-group contained both *stx_2b_+* and *stx-* isolates. However, the draft status of the genomes hindered further investigation of the ΦStx status.

Except for the H12 STEC, the LEE island is present in all other identified STEC phylogroups (H2, H6, H12, and H16) (**Fig. 7**). The LEE island is also present enteropathogenic *E. coli* (EPEC) (H5) and a strain containing genes associated with both enterotoxigenic *E. coli* (ETEC) and EPEC (H8) (**Fig. 7**). Over 20 intimins mediating the attachment to intestinal cells are currently described (58, 202–206). Previous studies indicated a correlation between the H-antigen and the intimin subtype in LEE-positive *E. coli* (207–209). Indeed, we found a correlation between H-antigens and the five *eae-*suballeles present in serogroup O118. The intimin *ε*-subtype is associated with H2 STEC, exemplified by strain EC20017429, and LST strains of this group (**Fig. 7**). This group also contains *eae*-negative strain PNUSAE161722, missing integral LEE components and effectors (**Fig. 7**). Its position in the tree and intimate relationship to ST-17 LEE+ isolates may suggest secondary loss rather than an ancestral LEE-status. The H16-phylogroup features the *β*-intimin, represented by strains 2014C-3050, 2013C-4538, 12867, and 12089 (**Fig. 4**). We further observed the following correlations H6-*eae-ι*, H5-*eae*-*κ*, and H8-*eae*-*θ*, in the other LEE+ phylogroups, though these were represented by only a few strains.

### Antimicrobial– and multidrug-resistance loci in serogroup O118

Antimicrobial-resistant and multidrug-resistant (MDR) *E. coli* O118 STEC strains have been previously reported (55, 124, 210). Consistent with these observations, AR/MDR strains were detected within STEC phylogroups H16 and H2 (Fig. 8), and not in other STEC lineages. Among the resistant O118 STEC, we identified genes conferring resistance to ten antibiotic classes: β-Lactam, Chloramphenicol (CHL), Quinolone (QNL), Tetracycline (Tet), Polymyxin (PMX), Macrolide–Lincosamide–Streptogramin (MLS), Aminoglycoside, Sulfonamide, Trimethoprim, and Rifampin (RIF). Numerous MDR isolates, as defined by resistance to three or more antibiotic classes, were detected. A prominent three-class profile— tetracyclines, aminoglycosides, and sulfonamides—was observed frequently in H16 and H2 STEC, similar to the one observed in the MDRVC-Island of strain 12089 (H16, ST21) (**Fig. 5**). We also noted that resistance gene presence and combinations followed phylogroup-specific patterns, as evident from the cgMLST-based phylogeny (Fig. 8). However, due to the draft status of most genomes, we were limited in our ability to determine the genomic context, whether these AR loci are chromosomal (**Fig. 5**), plasmid-borne (124) or phage-associated (13) (**Fig. 8**).

**FIG 8.**
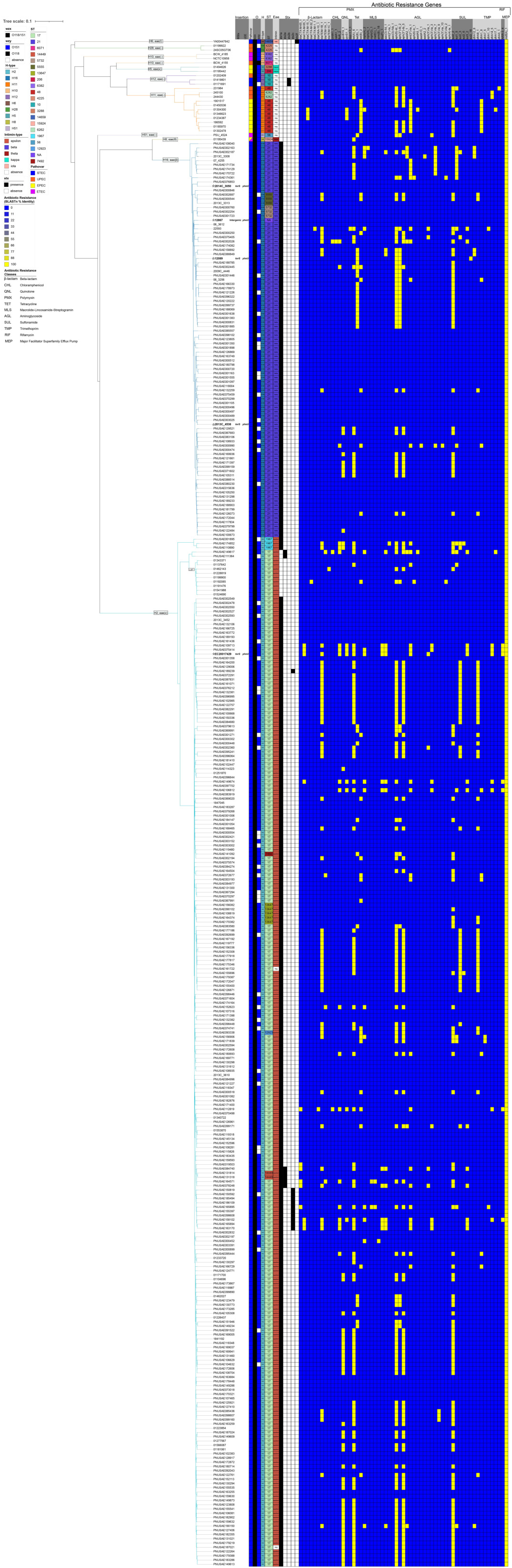
Phylogeny and distribution of antibiotic-resistance genes. Percentage identities for each antibiotic-resistance gene identified in ResFinder are visualized in a heatmap. The strains feature a total of 58 antibiotic-resistance genes that were classified into 10 antibiotic classes.

## Conclusion

Whole-genome sequencing (WGS) typing strategies have significantly advanced our understanding of the genomic makeup and evolution of *E. coli* pathovars (12, 211, 212). The high-resolution framework established for the *E. coli* serogroup O118/O151 complex allowed us to define the phylogenomic boundaries and virulence traits of emerging O118 STEC. For partitioning O118 serogroup isolates, their H-antigen was a robust phylogenetic marker (213), although genetically unrelated *E. coli* strains may share H-types due to lateral acquisition (214). Among the delineated STEC O118 phylogroups, we observed substantial plasticity in virulence and resistance inventories, characterized by diverse *stx* and *eae* suballeles. The presence or absence of *stx* was shaped by dynamic ΦStx prophage acquisitions but did not necessarily reflect evolutionary relationships (201). Notably, we identified a secondary Stx prophage loss event in an *stx*-negative H2 subgroup branching from ST-17 STEC, suggesting that such losses, followed by potential reacquisition, may lead to transitional shifts in pathogenic potential (125). Our findings showed a strict correlation between the H-antigen and LEE-borne intimin subtype in STEC (H6/*eae-ι*, H16/*eae-β*, H2/*eae-ε*) and non-STEC (H5/*eae-κ*, H8/*eae-θ*) isolates, reinforcing the hypothesis of evolutionary lineage sorting (207, 215–217). We identified horizontally acquired pathogenicity-associated islands (PAI) that enhance virulence potential. Notable was the O-island 122 inserted into the LEE-island of clinical H2-strain EC20017429, a locus previously associated with severe disease outcomes (173–175). The discovered multifunctional island in clinical H16-strain 12089 enhances interbacterial competitiveness, virulence, stress adaptation, and may increase the strain’s fitness and pathogenic potential. However, disease outcome is multifactorial, resulting from the complex interplay between the infective agent, the host microbiota (218–221), and the infected individual (222–225), and thus cannot be reliably predicted *in silico*. The emergence of clinically relevant O118 STEC in both human and animal reservoirs, with the potential to cause severe foodborne disease, highlights the need for ongoing genomic surveillance. Insights into pathogenome plasticity and evolutionary trajectories, gained through comparative genomic approaches, are critical for informing public health strategies, risk assessment, and infection control efforts.

## Supplemental Figures

**FIG S1 Phylogenetic relationship inferred from Average Nucleotide Identities.** UPGMA-clustered distance matrix of ANI values computed by an all-vs-all comparison of the 359 genomes.

**FIG S2 IS elements in the closed STEC O118 chromosomes.** This heatmap shows the prevalence and distribution of the 726 cataloged IS elements closed H2 and H16 O118 STEC, which can be categorized into 16 known families and 40 clusters. As determined by MLST typing, individual strain relationships are also reflected in IS cluster types and copy numbers.

**FIG S3 Comparison of closed O118 genome architectures.** Mauve-comparisons of chromosomes highlighting genomic rearrangements associated with hot spots for recombination.

**FIG S4 Pangenome analysis of serogroup O118**. The pangenome was computed with Roary, and the resulting gene presence/absence matrix was visualized in Phandango. The guide tree reflects the cgMLST tree topology, which partitions the isolates by H-type.

**FIG S5 Comparison of the ΦStx1a-prophages insertion sites in H2 strains.** BLASTn-based comparison of the ΦStx_1a_ prophage insertion sites in H2 stx+/-isolates. The comparisons show that phage remnants were detected in related *stx_1a_* negative phylogroups (H2, *stx-*) with the same phage integrase type at *torS*.

## Supplemental Tables

**TABLE S1** Strain-associated metadata and genome statistics

**TABLE S2** Predicted prophage and mobilome gene content

**TABLE S3** Predicted virulence and antimicrobial resistance gene content

**TABLE S4** Sequence Types according to the Achtman MLST schema

## Supplemental Text

### Plasmid profiles of serogroup O118 *E. coli*

The 359 analyzed closed and draft genomes identified 36 replicons, which provided a testament to the plasticity of plasmids in this group. Among these were incompatibility group families associated with virulence plasmids and colicins (**Table S2-6**). The most prevalent replicon in STEC O118 was IncFIB(AP001918) of H2 strains (97.1%, 235/242) and of H16 strains (94.6%, 88/93). These usually 100 kb or larger low-copy plasmids have been associated with disseminating virulence genes in *Enterobacteriaceae* (226). Other Inc-types with shared distribution were colicin type Col156, present in H2 (42.6%, 103/242) and H16 (7.5%, 7/93) strains, as well as present in the pO111 plasmid in H16 (29.0%, 27/93) and H2 (14.5%, 35/242) strains. The presence of Col156 is often associated with plasmids that also carry antibiotic resistance genes (227). Carriage of virulence plasmid pO111 has been linked to severe gastrointestinal disease (228). Another noteworthy observation was that most STEC H16– and H2-phylogroups feature IncB/O/K/Z_3 groups (95.7% and 90.7%), and IncFII(pCoo) that is present in all H11-strains.

## Data Availability Statement

The sequence data sets generated and analyzed in this study have been deposited in the Sequence Read Archive (SRA) and GenBank at NCBI under BioProject PRJNA1103183 (https://www.ncbi.nlm.nih.gov/search/all/?term=PRJNA1103183). Accessions for genomic reads, assembled annotated chromosomes, and plasmids, along with strain-associated metadata, are provided in **Table S1.**

## Author Contributions

Conceived and designed the experiments: JMB and ME. Analyzed the data: IR, SSK, JMB, and ME. Drafted the manuscript: ME. Contributed to and edited the manuscript: IR, SKK, JMB, and ME. All authors have read and agreed to the published version of this manuscript.

## Acknowledgments

This work received computational support from the High-Performance Computing Cluster (HPCC) operated by Tech Solutions at UTSA. The use of product and company names is necessary to accurately report the methods and results; however, the United States Department of Agriculture (USDA) neither guarantees nor warrants the standard of the products, and the use of names by the USDA implies no approval of the product to the exclusion of others that may also be suitable. The USDA is an equal opportunity provider and employer. We wish to thank Drs. Peter Iwen and Paul Fey of the University of Nebraska Medical Center College of Medicine and the Nebraska State Health Laboratory for providing *E. coli* strains 12089 and 12867. We would like to acknowledge Mariana Sainz Garcia, Felix Borrego, and Jacob Alford for assistance with data visualization.

## Bibliography

1. Stordeur P, China B, Charlier G, Roels S, Mainil J. 2000. Clinical signs, reproduction of attaching/effacing lesions, and enterocyte invasion after oral inoculation of an O118 enterohaemorrhagic Escherichia coli in neonatal calves. Microbes Infect 2:17–24.

2. Karmali MA, Petric M, Lim C, Fleming PC, Steele BT. 1983. Escherichia coli cytotoxin, haemolytic-uraemic syndrome, and haemorrhagic colitis. Lancet 2:1299–1300.

3. Majowicz SE, Scallan E, Jones-Bitton A, Sargeant JM, Stapleton J, Angulo FJ, Yeung DH, Kirk MD. 2014. Global incidence of human Shiga toxin-producing Escherichia coli infections and deaths: a systematic review and knowledge synthesis. Foodborne Pathog Dis 11:447–55.

4. Beutin L, Bulte M, Weber A, Zimmermann S, Gleier K. 2000. Investigation of human infections with verocytotoxin-producing strains of Escherichia coli (VTEC) belonging to serogroup O118 with evidence for zoonotic transmission. Epidemiol Infect 125:47–54.

5. Abu-Ali GS, Lacher DW, Wick LM, Qi W, Whittam TS. 2009. Genomic diversity of pathogenic Escherichia coli of the EHEC 2 clonal complex. BMC Genomics 10:296.

6. Maria Ferreira Cavalcanti A, Tavanelli Hernandes R, Harummyy Takagi E, Ernestina Cabílio Guth B, de Lima Ori É, Regina Schicariol Pinheiro S, Sueli de Andrade T, Louzada Oliveira S, Cecilia Cergole-Novella M, Rodrigues Francisco G, dos Santos LF. 2020. Virulence Profiling and Molecular Typing of Shiga Toxin-Producing E. Coli (STEC) From Human Sources in Brazil. 8:171.

7. Pearce MC, Evans J, McKendrick IJ, Smith AW, Knight HI, Mellor DJ, Woolhouse ME, Gunn GJ, Low JC. 2006. Prevalence and virulence factors of Escherichia coli serogroups O26, O103, O111, and O145 shed by cattle in Scotland. Appl Environ Microbiol 72:653–9.

8. Nataro JP, Kaper JB. 1998. Diarrheagenic Escherichia coli. Clin Microbiol Rev 11:142–201.

9. Kaper JB, Nataro JP, Mobley HL. 2004. Pathogenic Escherichia coli. Nat Rev Microbiol 2:123–40.

10. Basavaraju M, B. S G. 2022. Escherichia coli: An Overview of Main Characteristics, p 21 doi:10.5772/intechopen.105508.

11. Sanjar F, Hazen TH, Shah SM, Koenig SS, Agrawal S, Daugherty S, Sadzewicz L, Tallon LJ, Mammel MK, Feng P, Soderlund R, Tarr PI, Debroy C, Dudley EG, Cebula TA, Ravel J, Fraser CM, Rasko DA, Eppinger M. 2014. Genome Sequence of Escherichia coli O157:H7 Strain 2886-75, Associated with the First Reported Case of Human Infection in the United States. Genome Announc 2.

12. Rusconi B, Sanjar F, Koenig SS, Mammel MK, Tarr PI, Eppinger M. 2016. Whole Genome Sequencing for Genomics-Guided Investigations of Escherichia coli O157:H7 Outbreaks. Front Microbiol 7:985.

13. Eppinger M, Mammel MK, Leclerc JE, Ravel J, Cebula TA. 2011. Genomic anatomy of Escherichia coli O157:H7 outbreaks. Proc Natl Acad Sci U S A 108:20142–7.

14. Eppinger M, Daugherty S, Agrawal S, Galens K, Sengamalay N, Sadzewicz L, Tallon L, Cebula TA, Mammel MK, Feng P, Soderlund R, Tarr PI, Debroy C, Dudley EG, Fraser CM, Ravel J. 2013. Whole-Genome Draft Sequences of 26 Enterohemorrhagic Escherichia coli O157:H7 Strains. Genome Announc 1:e0013412.

15. Riley LW, Remis RS, Helgerson SD, McGee HB, Wells JG, Davis BR, Hebert RJ, Olcott ES, Johnson LM, Hargrett NT, Blake PA, Cohen ML. 1983. Hemorrhagic colitis associated with a rare Escherichia coli serotype. N Engl J Med 308:681–5.

16. Glassman H, Ferrato C, Chui L. 2022. Epidemiology of Non-O157 Shiga Toxin-Producing Escherichia coli in the Province of Alberta, Canada, from 2018 to 2021. Microorganisms 10.

17. Vishram B, Jenkins C, Greig DR, Godbole G, Carroll K, Balasegaram S, Byrne L. 2021. The emerging importance of Shiga toxin-producing Escherichia coli other than serogroup O157 in England. J Med Microbiol 70.

18. Gould LH, Mody RK, Ong KL, Clogher P, Cronquist AB, Garman KN, Lathrop S, Medus C, Spina NL, Webb TH, White PL, Wymore K, Gierke RE, Mahon BE, Griffin PM. 2013. Increased recognition of non-O157 Shiga toxin-producing Escherichia coli infections in the United States during 2000-2010: epidemiologic features and comparison with E. coli O157 infections. Foodborne Pathog Dis 10:453–60.

19. Tarr GAM, Rounds J, Vachon MS, Smith K, Medus C, Hedberg CW. 2023. Differences in risk factors for transmission among Shiga toxin-producing Escherichia coli serogroups and stx profiles. J Infect 87:498–505.

20. Kalalah AA, Koenig SSK, Bono JL, Bosilevac JM, Eppinger M. 2024. Pathogenomes and virulence profiles of representative big six non-O157 serogroup Shiga toxin-producing Escherichia coli. Front Microbiol 15:1364026.

21. (CDC) CfDCaP. 2021. National Shiga toxin-producing Escherichia coli (STEC) Surveillance Annual Report, 2017. Atlanta GUDoHaHS,

22. Tseng M, Sha Q, Rudrik JT, Collins J, Henderson T, Funk JA, Manning SD. 2016. Increasing incidence of non-O157 Shiga toxin-producing Escherichia coli (STEC) in Michigan and association with clinical illness. Epidemiol Infect 144:1394–405.

23. Blankenship HM, Mosci RE, Phan Q, Fontana J, Rudrik JT, Manning SD. 2020. Genetic Diversity of Non-O157 Shiga Toxin-Producing Escherichia coli Recovered From Patients in Michigan and Connecticut. Front Microbiol 11:529.

24. Luna-Gierke RE, Griffin PM, Gould LH, Herman K, Bopp CA, Strockbine N, Mody RK. 2014. Outbreaks of non-O157 Shiga toxin-producing Escherichia coli infection: USA. Epidemiol Infect 142:2270–80.

25. Blankenship HM, Mosci RE, Dietrich S, Burgess E, Wholehan J, McWilliams K, Pietrzen K, Benko S, Gatesy T, Rudrik JT, Soehnlen M, Manning SD. 2021. Population structure and genetic diversity of non-O157 Shiga toxin-producing Escherichia coli (STEC) clinical isolates from Michigan. Sci Rep 11:4461.

26. Gould LH, Mody RK, Ong KL, Clogher P, Cronquist AB, Garman KN, Lathrop S, Medus C, Spina NL, Webb TH, White PL, Wymore K, Gierke RE, Mahon BE, Griffin PM, Emerging Infections Program Foodnet Working G. 2013. Increased recognition of non-O157 Shiga toxin-producing Escherichia coli infections in the United States during 2000-2010: epidemiologic features and comparison with E. coli O157 infections. Foodborne Pathog Dis 10:453–60.

27. Alharbi MG, Al-Hindi RR, Esmael A, Alotibi IA, Azhari SA, Alseghayer MS, Teklemariam AD. 2022. The “Big Six”: Hidden Emerging Foodborne Bacterial Pathogens. Tropical Medicine and Infectious Disease 7:356.

28. Hashimoto H, Mizukoshi K, Nishi M, Kawakita T, Hasui S, Kato Y, Ueno Y, Takeya R, Okuda N, Takeda T. 1999. Epidemic of gastrointestinal tract infection including hemorrhagic colitis attributable to Shiga toxin 1-producing Escherichia coli O118:H2 at a junior high school in Japan. Pediatrics 103:E2.

29. Wieler LH, Busse B, Steinruck H, Beutin L, Weber A, Karch H, Baljer G. 2000. Enterohemorrhagic Escherichia coli (EHEC) strains of serogroup O118 display three distinctive clonal groups of EHEC pathogens. J Clin Microbiol 38:2162–9.

30. Orskov F, Orskov I, Furowicz AJ. 1972. Four new Escherichia coli O antigens, O148, O151, O152, O153, and one new H antigen, H50, found in strains isolated from enteric diseases in man with a discussion on the future numbering of K antigens. Acta Pathol Microbiol Scand B Microbiol Immunol 80:435–40.

31. Steinruck H, Mochmann H, Kontny I, Werner U. 1980. [Escherichia coli O151:K-:h10 (“krim”) as etiologic agent of diarrhoea (author’s transl)]. Zentralbl Bakteriol A 248:335–44.

32. Wieler LH, Schwanitz A, Vieler E, Busse B, Steinruck H, Kaper JB, Baljer G. 1998. Virulence properties of Shiga toxin-producing Escherichia coli (STEC) strains of serogroup O118, a major group of STEC pathogens in calves. J Clin Microbiol 36:1604–7.

33. Castro VS, Figueiredo EES, Stanford K, McAllister T, Conte-Junior CA. 2019. Shiga-Toxin Producing Escherichia Coli in Brazil: A Systematic Review. Microorganisms 7.

34. Kruger A, Lucchesi PM. 2015. Shiga toxins and stx phages: highly diverse entities. Microbiology (Reading) 161:451–62.

35. Skinner C, Patfield S, Stanker LH, Fratamico P, He X. 2014. New high-affinity monoclonal antibodies against Shiga toxin 1 facilitate the detection of hybrid Stx1/Stx2 in vivo. PLoS One 9:e99854.

36. Sperandio V, Hovde CJ. 2015. Enterohemorrhagic Escherichia coli and other Shiga toxin-producing E. coli ASM Press, Washington, DC.

37. Zuppi M, Tozzoli R, Chiani P, Quiros P, Martinez-Velazquez A, Michelacci V, Muniesa M, Morabito S. 2020. Investigation on the Evolution of Shiga Toxin-Converting Phages Based on Whole Genome Sequencing. Front Microbiol 11:1472.

38. Asadulghani M, Ogura Y, Ooka T, Itoh T, Sawaguchi A, Iguchi A, Nakayama K, Hayashi T. 2009. The defective prophage pool of Escherichia coli O157: prophage-prophage interactions potentiate horizontal transfer of virulence determinants. PLoS Pathog 5:e1000408.

39. Huang A, Friesen J, Brunton JL. 1987. Characterization of a bacteriophage that carries the genes for production of Shiga-like toxin 1 in Escherichia coli. J Bacteriol 169:4308–12.

40. Alizade H, Hosseini Teshnizi S, Azad M, Shojae S, Gouklani H, Davoodian P, Ghanbarpour R. 2019. An overview of diarrheagenic Escherichia coli in Iran: A systematic review and meta-analysis. J Res Med Sci 24:23.

41. Bielaszewska M, Mellmann A, Zhang W, Kock R, Fruth A, Bauwens A, Peters G, Karch H. 2011. Characterisation of the Escherichia coli strain associated with an outbreak of haemolytic uraemic syndrome in Germany, 2011: a microbiological study. Lancet Infect Dis 11:671–6.

42. Beutin L, Miko A, Krause G, Pries K, Haby S, Steege K, Albrecht N. 2007. Identification of human-pathogenic strains of Shiga toxin-producing Escherichia coli from food by a combination of serotyping and molecular typing of Shiga toxin genes. Appl Environ Microbiol 73:4769–75.

43. Hauser JR, Atitkar RR, Petro CD, Lindsey RL, Strockbine N, O’Brien AD, Melton-Celsa AR. 2020. The Virulence of Escherichia coli O157:H7 Isolates in Mice Depends on Shiga Toxin Type 2a (Stx2a)-Induction and High Levels of Stx2a in Stool. Front Cell Infect Microbiol 10:62.

44. Fuller CA, Pellino CA, Flagler MJ, Strasser JE, Weiss AA. 2011. Shiga toxin subtypes display dramatic differences in potency. Infect Immun 79:1329–37.

45. Pinto G, Sampaio M, Dias O, Almeida C, Azeredo J, Oliveira H. 2021. Insights into the genome architecture and evolution of Shiga toxin encoding bacteriophages of Escherichia coli. BMC Genomics 22:366.

46. McNichol BA, Bova RA, Torres K, Preston LN, Melton-Celsa AR. 2021. Switching Shiga Toxin (Stx) Type from Stx2d to Stx2a but Not Stx2c Alters Virulence of Stx-Producing Escherichia coli (STEC) Strain B2F1 in Streptomycin (Str)-Treated Mice. Toxins 13:64.

47. Bai X, Zhang W, Tang X, Xin Y, Xu Y, Sun H, Luo X, Pu J, Xu J, Xiong Y, Lu S. 2016. Shiga Toxin-Producing Escherichia coli in Plateau Pika (Ochotona curzoniae) on the Qinghai-Tibetan Plateau, China. Front Microbiol 7:375.

48. Usein CR, Ciontea AS, Militaru CM, Condei M, Dinu S, Oprea M, Cristea D, Michelacci V, Scavia G, Zota LC, Zaharia A, Morabito S. 2017. Molecular characterisation of human Shiga toxin-producing Escherichia coli O26 strains: results of an outbreak investigation, Romania, February to August 2016. Euro Surveill 22.

49. Yamasaki E, Watahiki M, Isobe J, Sata T, Nair G, Kurazono H. 2015. Quantitative Detection of Shiga Toxins Directly From Stool Specimens of Patients Associated With an Outbreak of Enterohemorrhagic Escherichia Coli in Japan—Quantitative Shiga Toxin Detection From Stool During EHEC Outbreak. 7:4381–4389.

50. Petro CD, Trojnar E, Sinclair J, Liu Z-M, Smith M, O’Brien AD, Melton-Celsa A. 2019. Shiga Toxin Type 1a (Stx1a) Reduces the Toxicity of the More Potent Stx2a In Vivo and In Vitro. 87.

51. Liu DL, Cui GH, Cai JP. 2008. [Surveillance on E. coli O157: H7 in Dongtai in Jiangsu province]. Zhonghua Liu Xing Bing Xue Za Zhi 29:202.

52. Franzin FM, Sircili MP. 2015. Locus of enterocyte effacement: a pathogenicity island involved in the virulence of enteropathogenic and enterohemorragic Escherichia coli subjected to a complex network of gene regulation. Biomed Res Int 2015:534738.

53. Dutta S, Pazhani GP, Nataro JP, Ramamurthy T. 2015. Heterogenic Virulence in a Diarrheagenic Escherichia Coli: Evidence for an EPEC Expressing Heat-Labile Toxin of ETEC. International Journal of Medical Microbiology 305:47–54.

54. Carrilero L, Dunn SJ, Moran RA, McNally A, Brockhurst MA. 2023. Evolutionary Responses to Acquiring a Multidrug Resistance Plasmid Are Dominated by Metabolic Functions across Diverse Escherichia coli Lineages. mSystems 8:e0071322.

55. Maidhof H, Guerra B, Abbas S, Elsheikha HM, Whittam TS, Beutin L. 2002. A multiresistant clone of Shiga toxin-producing Escherichia coli O118:[H16] is spread in cattle and humans over different European countries. Appl Environ Microbiol 68:5834–42.

56. Ogura Y, Ooka T, Iguchi A, Toh H, Asadulghani M, Oshima K, Kodama T, Abe H, Nakayama K, Kurokawa K, Tobe T, Hattori M, Hayashi T. 2009. Comparative genomics reveal the mechanism of the parallel evolution of O157 and non-O157 enterohemorrhagic Escherichia coli. Proc Natl Acad Sci U S A 106:17939–44.

57. Caprioli A, Morabito S, Brugère H, Oswald E. 2005. Enterohaemorrhagic Escherichia coli: emerging issues on virulence and modes of transmission. Vet Res 36:289–311.

58. Kalalah AA, Koenig SSK, Bono JL, Bosilevac JM, Eppinger M. 2024. Pathogenomes and virulence profiles of representative big six non-O157 serogroup Shiga toxin-producing Escherichia coli. Frontiers in Microbiology 15.

59. Boisen N, Melton-Celsa AR, Hansen AM, Zangari T, Smith MA, Russo LM, Scheutz F, O’Brien AD, Nataro JP. 2019. The Role of the AggR Regulon in the Virulence of the Shiga Toxin-Producing Enteroaggregative Escherichia coli Epidemic O104:H4 Strain in Mice. Front Microbiol 10:1824.

60. Boisen N, Melton-Celsa AR, Scheutz F, O’Brien AD, Nataro JP. 2015. Shiga toxin 2a and Enteroaggregative Escherichia coli––a deadly combination. Gut Microbes 6:272–8.

61. Van Nederveen V, Melton-Celsa A. 2024. Extracellular components in enteroaggregative Escherichia coli biofilm and impact of treatment with proteinase K, DNase or sodium metaperiodate. Front Cell Infect Microbiol 14:1379206.

62. Liu Y, Gilchrist A, Zhang J, Li XF. 2008. Detection of viable but nonculturable Escherichia coli O157:H7 bacteria in drinking water and river water. Appl Environ Microbiol 74:1502–7.

63. Community TG. 2022. The Galaxy platform for accessible, reproducible and collaborative biomedical analyses: 2022 update. Nucleic Acids Research 50:W345–W351.

64. Kolmogorov M, Yuan J, Lin Y, Pevzner PA. 2019. Assembly of long, error-prone reads using repeat graphs. Nature Biotechnology 37:540–546.

65. Tatusova T, DiCuccio M, Badretdin A, Chetvernin V, Nawrocki EP, Zaslavsky L, Lomsadze A, Pruitt KD, Borodovsky M, Ostell J. 2016. NCBI prokaryotic genome annotation pipeline. Nucleic Acids Res 44:6614–24.

66. Shen W, Sipos B, Zhao L. 2024. SeqKit2: A Swiss army knife for sequence and alignment processing. Imeta 3:e191.

67. Shen W, Le S, Li Y, Hu F. 2016. SeqKit: A Cross-Platform and Ultrafast Toolkit for FASTA/Q File Manipulation. PLoS One 11:e0163962.

68. Blattner FR, Plunkett G, 3rd, Bloch CA, Perna NT, Burland V, Riley M, Collado-Vides J, Glasner JD, Rode CK, Mayhew GF, Gregor J, Davis NW, Kirkpatrick HA, Goeden MA, Rose DJ, Mau B, Shao Y. 1997. The complete genome sequence of Escherichia coli K-12. Science 277:1453–62.

69. Riley M, Abe T, Arnaud MB, Berlyn MK, Blattner FR, Chaudhuri RR, Glasner JD, Horiuchi T, Keseler IM, Kosuge T, Mori H, Perna NT, Plunkett G, 3rd, Rudd KE, Serres MH, Thomas GH, Thomson NR, Wishart D, Wanner BL. 2006. Escherichia coli K-12: a cooperatively developed annotation snapshot––2005. Nucleic Acids Res 34:1–9.

70. Jünemann S, Sedlazeck FJ, Prior K, Albersmeier A, John U, Kalinowski J, Mellmann A, Goesmann A, von Haeseler A, Stoye J, Harmsen D. 2013. Updating benchtop sequencing performance comparison. Nature Biotechnology 31:294–296.

71. Foley SL, Lynne AM, Nayak R. 2009. Molecular typing methodologies for microbial source tracking and epidemiological investigations of Gram-negative bacterial foodborne pathogens. Infect Genet Evol 9:430–40.

72. Zhou Z, Alikhan NF, Mohamed K, Fan Y, Agama Study G, Achtman M. 2019. The EnteroBase user’s guide, with case studies on Salmonella transmissions, Yersinia pestis phylogeny and Escherichia core genomic diversity. Genome Res doi:10.1101/gr.251678.119.

73. Zhou Z, Alikhan NF, Mohamed K, Fan Y, Achtman M. 2020. The EnteroBase user’s guide, with case studies on Salmonella transmissions, Yersinia pestis phylogeny, and Escherichia core genomic diversity. Genome Res 30:138–152.

74. Díaz L, Gutierrez S, Moreno-Switt AI, Hervé LP, Hamilton-West C, Padola NL, Navarrete P, Reyes-Jara A, Meng J, González-Escalona N, Toro M. 2021. Diversity of Non-O157 Shiga Toxin-Producing Escherichia coli Isolated from Cattle from Central and Southern Chile. Animals (Basel) 11.

75. Kruskal JB. 1956. On the Shortest Spanning Subtree of a Graph and the Traveling Salesman Problem. Proceedings of the American Mathematical Society 7:48–50.

76. Francisco AP, Bugalho M, Ramirez M, Carriço JA. 2009. Global optimal eBURST analysis of multilocus typing data using a graphic matroid approach. BMC Bioinformatics 10:152.

77. Darling AE, Mau B, Perna NT. 2010. progressiveMauve: Multiple Genome Alignment with Gene Gain, Loss and Rearrangement. PloS one 5:e11147.

78. Shimoyama Y. 2024. pyGenomeViz: A genome visualization python package for comparative genomics.

79. Waskom ML. 2021. seaborn: statistical data visualization. ournal of Open Source Software 6(60), 3021.

80. Alikhan NF, Petty NK, Ben Zakour NL, Beatson SA. 2011. BLAST Ring Image Generator (BRIG): simple prokaryote genome comparisons. BMC Genomics 12:402.

81. Shaw J, Yu YW. 2023. Fast and robust metagenomic sequence comparison through sparse chaining with skani. Nat Methods 20:1661–1665.

82. Liu B, Zheng D, Zhou S, Chen L, Yang J. 2022. VFDB 2022: a general classification scheme for bacterial virulence factors. Nucleic Acids Res 50:D912–D917.

83. Florensa AF, Kaas RS, Clausen P, Aytan-Aktug D, Aarestrup FM. 2022. ResFinder – an open online resource for identification of antimicrobial resistance genes in next-generation sequencing data and prediction of phenotypes from genotypes. Microb Genom 8.

84. Altschul SF, Gish W, Miller W, Myers EW, Lipman DJ. 1990. Basic local alignment search tool. J Mol Biol 215:403–10.

85. Baral SK. 2024. Characterization of Virulence Factors in Multidrug-Resistant Escherichia Coli Isolated From Intestinal and Extra – Intestinal Clinical Samples. Journal of Manmohan Memorial Institute of Health Sciences 9:13–18.

86. Boroumand M, Sharifi A, Ghatei MA, Sadrinasab M. 2021. Evaluation of Biofilm Formation and Virulence Genes and Association With Antibiotic Resistance Patterns of Uropathogenic Escherichia Coli Strains in Southwestern Iran. Jundishapur Journal of Microbiology 14.

87. Boudjerda D, Lahouel M. 2022. Virulence and Antimicrobial Resistance of Escherichia Coli Isolated From Chicken Meat, Beef, and Raw Milk. Austral Journal of Veterinary Sciences 54:115–125.

88. Coura FM, Diniz AN, Oliveira CA, Lage AP, Lobato FCF, Heinemann MB, Silva ROS. 2018. Detection of Virulence Genes and the Phylogenetic Groups of Escherichia Coli Isolated From Dogs in Brazil. Ciência Rural 48.

89. Lindstedt B-A, Finton MD, Porcellato D, Brandal LT. 2018. High Frequency of Hybrid Escherichia Coli Strains With Combined Intestinal Pathogenic Escherichia Coli (IPEC) and Extraintestinal Pathogenic Escherichia Coli (ExPEC) Virulence Factors Isolated From Human Faecal Samples. BMC Infectious Diseases 18.

90. Otokunefor K, Onyemelukwe A, Agbake B, Owhoigwe N. 2022. Molecular Detection of Virulence and Resistance Markers in Escherichia Coli and Staphylococcus Aureus Isolated From Nigerian Currency. Journal of Life and Bio Sciences Research 3:23–26.

91. Vanstokstraeten R, Crombé F, Piérard D, A CM, Wybo I, D DG, Janssen T, Caljon B, Demuyser T. 2022. Molecular Characterization of Extraintestinal and Diarrheagenic *Escherichia Coli* Blood Isolates. Virulence 13:2032–2041.

92. Yim J-H, Kun-Ho S, Chon JW, Jeong D, Song KY. 2021. Status and Prospects of PCR Detection Methods for Diagnosing Pathogenic Escherichia Coli: A Review. Journal of Dairy Science and Biotechnology 39:51–62.

93. Seemann T. 2013. barrnap 0.7: rapid ribosomal RNA prediction.

94. Couvin D, Bernheim A, Toffano-Nioche C, Touchon M, Michalik J, Neron B, Rocha EPC, Vergnaud G, Gautheret D, Pourcel C. 2018. CRISPRCasFinder, an update of CRISRFinder, includes a portable version, enhanced performance and integrates search for Cas proteins. Nucleic Acids Res 46:W246–W251.

95. Grant JR, Enns E, Marinier E, Mandal A, Herman EK, Chen CY, Graham M, Van Domselaar G, Stothard P. 2023. Proksee: in-depth characterization and visualization of bacterial genomes. Nucleic Acids Res 51:W484–W492.

96. Tesson F, Herve A, Mordret E, Touchon M, d’Humieres C, Cury J, Bernheim A. 2022. Systematic and quantitative view of the antiviral arsenal of prokaryotes. Nat Commun 13:2561.

97. Sullivan MJ, Petty NK, Beatson SA. 2011. Easyfig: a genome comparison visualizer. Bioinformatics 27:1009–10.

98. Wishart DS, Han S, Saha S, Oler E, Peters H, Grant JR, Stothard P, Gautam V. 2023. PHASTEST: faster than PHASTER, better than PHAST. Nucleic Acids Res 51:W443–W450.

99. Ogura Y, Mondal SI, Islam MR, Mako T, Arisawa K, Katsura K, Ooka T, Gotoh Y, Murase K, Ohnishi M, Hayashi T. 2015. The Shiga toxin 2 production level in enterohemorrhagic Escherichia coli O157:H7 is correlated with the subtypes of toxin-encoding phage. Sci Rep 5:16663.

100. Llarena AK, Aspholm M, O’Sullivan K, Wêgrzyn G, Lindbäck T. 2021. Replication Region Analysis Reveals Non-lambdoid Shiga Toxin Converting Bacteriophages. Front Microbiol 12:640945.

101. Fagerlund A, Aspholm M, Węgrzyn G, Lindbäck T. 2022. High diversity in the regulatory region of Shiga toxin encoding bacteriophages. BMC Genomics 23:230.

102. Scheutz F, Teel LD, Beutin L, Pierard D, Buvens G, Karch H, Mellmann A, Caprioli A, Tozzoli R, Morabito S, Strockbine NA, Melton-Celsa AR, Sanchez M, Persson S, O’Brien AD. 2012. Multicenter evaluation of a sequence-based protocol for subtyping Shiga toxins and standardizing Stx nomenclature. J Clin Microbiol 50:2951–63.

103. Ashton PM, Perry N, Ellis R, Petrovska L, Wain J, Grant KA, Jenkins C, Dallman TJ. 2015. Insight into Shiga toxin genes encoded by Escherichia coli O157 from whole genome sequencing. Peerj 3.

104. Carrillo CD, Koziol AG, Mathews A, Goji N, Lambert D, Huszczynski G, Gauthier M, Amoako K, Blais BW. 2016. Comparative Evaluation of Genomic and Laboratory Approaches for Determination of Shiga Toxin Subtypes in Escherichia coli. J Food Prot 79:2078–2085.

105. Bertelli C, Laird MR, Williams KP, Simon Fraser University Research Computing G, Lau BY, Hoad G, Winsor GL, Brinkman FSL. 2017. IslandViewer 4: expanded prediction of genomic islands for larger-scale datasets. Nucleic Acids Res 45:W30–W35.

106. Bertelli C, Brinkman FSL. 2018. Improved genomic island predictions with IslandPath-DIMOB. Bioinformatics 34:2161–2167.

107. Bertelli C, Tilley KE, Brinkman FSL. 2018. Microbial genomic island discovery, visualization and analysis. Brief Bioinform doi:10.1093/bib/bby042.

108. Camacho C, Coulouris G, Avagyan V, Ma N, Papadopoulos J, Bealer K, Madden TL. 2009. BLAST+: architecture and applications. BMC Bioinformatics 10:421.

109. Ross K, Varani AM, Snesrud E, Huang H, Alvarenga DO, Zhang J, Wu C, McGann P, Chandler M. 2021. TnCentral: a Prokaryotic Transposable Element Database and Web Portal for Transposon Analysis. mBio 12:e0206021.

110. Siguier P, Perochon J, Lestrade L, Mahillon J, Chandler M. 2006. ISfinder: the reference centre for bacterial insertion sequences. Nucleic Acids Res 34:D32–6.

111. Liu M, Li X, Xie Y, Bi D, Sun J, Li J, Tai C, Deng Z, Ou HY. 2019. ICEberg 2.0: an updated database of bacterial integrative and conjugative elements. Nucleic Acids Res 47:D660–D665.

112. Xie Z, Tang H. 2017. ISEScan: automated identification of insertion sequence elements in prokaryotic genomes. Bioinformatics 33:3340–3347.

113. Robertson J, Nash JHE. 2018. MOB-suite: software tools for clustering, reconstruction and typing of plasmids from draft assemblies. Microb Genom 4.

114. Carattoli A, Zankari E, Garcia-Fernandez A, Voldby Larsen M, Lund O, Villa L, Moller Aarestrup F, Hasman H. 2014. In silico detection and typing of plasmids using PlasmidFinder and plasmid multilocus sequence typing. Antimicrob Agents Chemother 58:3895–903.

115. Galata V, Fehlmann T, Backes C, Keller A. 2019. PLSDB: a resource of complete bacterial plasmids. Nucleic Acids Res 47:D195–D202.

116. Ondov BD, Treangen TJ, Melsted P, Mallonee AB, Bergman NH, Koren S, Phillippy AM. 2016. Mash: fast genome and metagenome distance estimation using MinHash. Genome Biol 17:132.

117. van Heel AJ, de Jong A, Song C, Viel JH, Kok J, Kuipers OP. 2018. BAGEL4: a user-friendly web server to thoroughly mine RiPPs and bacteriocins. Nucleic Acids Res 46:W278–W281.

118. Brown CL, Mullet J, Hindi F, Stoll JE, Gupta S, Choi M, Keenum I, Vikesland P, Pruden A, Zhang L. 2022. mobileOG-db: a Manually Curated Database of Protein Families Mediating the Life Cycle of Bacterial Mobile Genetic Elements. Appl Environ Microbiol 88:e0099122.

119. Seemann T. 2014. Prokka: rapid prokaryotic genome annotation. Bioinformatics 30:2068–2069.

120. Page AJ, Cummins CA, Hunt M, Wong VK, Reuter S, Holden MTG, Fookes M, Falush D, Keane JA, Parkhill J. 2015. Roary: rapid large-scale prokaryote pan genome analysis. Bioinformatics 31:3691–3693.

121. Page AJ, Cummins CA, Hunt M, Wong VK, Reuter S, Holden MT, Fookes M, Falush D, Keane JA, Parkhill J. 2015. Roary: rapid large-scale prokaryote pan genome analysis. Bioinformatics 31:3691–3.

122. Hadfield J, Croucher NJ, Goater RJ, Abudahab K, Aanensen DM, Harris SR. 2017. Phandango: an interactive viewer for bacterial population genomics. Bioinformatics 34:292–293.

123. Patel PN, Lindsey RL, Garcia-Toledo L, Rowe LA, Batra D, Whitley SW, Drapeau D, Stoneburg D, Martin H, Juieng P. 2018. High-quality whole-genome sequences for 77 Shiga toxin-producing Escherichia coli strains generated with PacBio sequencing. Genome announcements 6:10.1128/genomea.00391-18.

124. Abdullah S, Almusallam A, Li M, Mahmood MS, Mushtaq MA, Eltai NO, Toleman MA, Mohsin M. 2023. Whole Genome-Based Genetic Insights Of *bla*_NDM_ Producing Clinical *E*. Coli Isolates in Hospital Settings of Pakistant. doi:10.1128/spectrum.00584-23.

125. Nyong EC, Zaia SR, Allue-Guardia A, Rodriguez AL, Irion-Byrd Z, Koenig SSK, Feng P, Bono JL, Eppinger M. 2020. Pathogenomes of Atypical Non-shigatoxigenic Escherichia coli NSF/SF O157:H7/NM: Comprehensive Phylogenomic Analysis Using Closed Genomes. Front Microbiol 11:619.

126. Allue-Guardia A, Koenig SSK, Martinez RA, Rodriguez AL, Bosilevac JM, Feng P, Eppinger M. 2022. Pathogenomes and variations in Shiga toxin production among geographically distinct clones of Escherichia coli O113:H21. Microb Genom 8.

127. Rasko DA, Rosovitz MJ, Myers GS, Mongodin EF, Fricke WF, Gajer P, Crabtree J, Sebaihia M, Thomson NR, Chaudhuri R, Henderson IR, Sperandio V, Ravel J. 2008. The pangenome structure of Escherichia coli: comparative genomic analysis of E. coli commensal and pathogenic isolates. J Bacteriol 190:6881–93.

128. Jain C, Rodriguez-R LM, Phillippy AM, Konstantinidis KT, Aluru S. 2018. High throughput ANI analysis of 90K prokaryotic genomes reveals clear species boundaries. Nature Communications 9:5114.

129. Ohnishi M, Kurokawa K, Hayashi T. 2001. Diversification of Escherichia coli genomes: are bacteriophages the major contributors? Trends Microbiol 9:481–5.

130. Kruger A, Lucchesi PM. 2015. Shiga toxins and stx phages: highly diverse entities. Microbiology 161:451–62.

131. Garcia-Aljaro C, Muniesa M, Jofre J, Blanch AR. 2009. Genotypic and phenotypic diversity among induced, stx2-carrying bacteriophages from environmental Escherichia coli strains. Appl Environ Microbiol 75:329–36.

132. Tozzoli R, Grande L, Michelacci V, Ranieri P, Maugliani A, Caprioli A, Morabito S. 2014. Shiga toxin-converting phages and the emergence of new pathogenic Escherichia coli: a world in motion. Front Cell Infect Microbiol 4:80.

133. Delannoy S, Mariani-Kurkdjian P, Webb HE, Bonacorsi S, Fach P. 2017. The Mobilome; A Major Contributor to Escherichia coli stx2-Positive O26:H11 Strains Intra-Serotype Diversity. Front Microbiol 8:1625.

134. Perna NT, Plunkett G, 3rd, Burland V, Mau B, Glasner JD, Rose DJ, Mayhew GF, Evans PS, Gregor J, Kirkpatrick HA, Posfai G, Hackett J, Klink S, Boutin A, Shao Y, Miller L, Grotbeck EJ, Davis NW, Lim A, Dimalanta ET, Potamousis KD, Apodaca J, Anantharaman TS, Lin J, Yen G, Schwartz DC, Welch RA, Blattner FR. 2001. Genome sequence of enterohaemorrhagic Escherichia coli O157:H7. Nature 409:529–33.

135. Kaper JB. 2005. Pathogenic Escherichia coli. Int J Med Microbiol 295:355–6.

136. Toro M, Rump LV, Cao G, Meng J, Brown EW, Gonzalez-Escalona N. 2015. Simultaneous presence of insertion sequence-excision enhancer (IEE) and insertion sequence IS629 correlates with increased diversity and virulence in Shiga-toxin producing Escherichia coli (STEC). J Clin Microbiol doi:10.1128/JCM.01349-15.

137. Stanton E, Park D, Dopfer D, Ivanek R, Kaspar CW. 2014. Phylogenetic characterization of Escherichia coli O157: H7 based on IS629 distribution and Shiga toxin genotype. Microbiology 160:502–13.

138. Yokoyama E, Hashimoto R, Etoh Y, Ichihara S, Horikawa K, Uchimura M. 2011. Biased distribution of IS629 among strains in different lineages of enterohemorrhagic Escherichia coli serovar O157. Infect Genet Evol 11:78–82.

139. Frost LS, Leplae R, Summers AO, Toussaint A. 2005. Mobile genetic elements: the agents of open source evolution. Nature Reviews Microbiology 3:722–732.

140. Hacker J, Blum-Oehler G, Hochhut B, Dobrindt U. 2003. The molecular basis of infectious diseases: pathogenicity islands and other mobile genetic elements. A review. Acta Microbiol Immunol Hung 50:321–30.

141. Dobrindt U, Blum-Oehler G, Nagy G, Schneider G, Johann A, Gottschalk G, Hacker J. 2002. Genetic structure and distribution of four pathogenicity islands (PAI I(536) to PAI IV(536)) of uropathogenic Escherichia coli strain 536. Infect Immun 70:6365–72.

142. Hashimoto JG, Stevenson BS, Schmidt TM. 2003. Rates and consequences of recombination between rRNA operons. J Bacteriol 185:966–72.

143. Cascales E, Buchanan SK, Duche D, Kleanthous C, Lloubes R, Postle K, Riley M, Slatin S, Cavard D. 2007. Colicin biology. Microbiol Mol Biol Rev 71:158–229.

144. Masaki H, Ogawa T. 2002. The modes of action of colicins E5 and D, and related cytotoxic tRNases. Biochimie 84:433–8.

145. Budič M, Rijavec M, Petkovšek Ž, Žgur-Bertok D. 2011. Escherichia Coli Bacteriocins: Antimicrobial Efficacy and Prevalence Among Isolates From Patients With Bacteraemia. Plos One 6:e28769.

146. Petrova V, Chitteni-Pattu S, Drees JC, Inman RB, Cox MM. 2009. An SOS inhibitor that binds to free RecA protein: the PsiB protein. Mol Cell 36:121–30.

147. Johnson TJ, Johnson SJ, Nolan LK. 2006. Complete DNA sequence of a ColBM plasmid from avian pathogenic Escherichia coli suggests that it evolved from closely related ColV virulence plasmids. J Bacteriol 188:5975–83.

148. Gigliucci F, van Hoek A, Chiani P, Knijn A, Minelli F, Scavia G, Franz E, Morabito S, Michelacci V. 2021. Genomic Characterization of hlyF-positive Shiga Toxin-Producing Escherichia coli, Italy and the Netherlands, 2000-2019. Emerg Infect Dis 27:853–861.

149. Tontanahal A, Sperandio V, Kovbasnjuk O, Loos S, Kristoffersson AC, Karpman D, Arvidsson I. 2022. IgG Binds Escherichia coli Serine Protease EspP and Protects Mice From E. coli O157:H7 Infection. Front Immunol 13:807959.

150. Yin X, Zhu J, Feng Y, Chambers JR, Gong J, Gyles CL. 2011. Differential gene expression and adherence of Escherichia coli O157:H7 in vitro and in ligated pig intestines. PLoS One 6:e17424.

151. Abdel-Shafi S, El-Serwy H, El-Zawahry Y, Zaki M, Sitohy B, Sitohy M. 2022. The Association between icaA and icaB Genes, Antibiotic Resistance and Biofilm Formation in Clinical Isolates of Staphylococci spp. Antibiotics (Basel) 11.

152. Anisimov AP, Shaikhutdinova RZ, Pan’kina LN, Feodorova VA, Savostina EP, Bystrova OV, Lindner B, Mokrievich AN, Bakhteeva IV, Titareva GM, Dentovskaya SV, Kocharova NA, Senchenkova SN, Holst O, Devdariani ZL, Popov YA, Pier GB, Knirel YA. 2007. Effect of deletion of the lpxM gene on virulence and vaccine potential of Yersinia pestis in mice. J Med Microbiol 56:443–453.

153. Frost LS, Leplae R, Summers AO, Toussaint A. 2005. Mobile genetic elements: the agents of open source evolution. Nat Rev Microbiol 3:722–32.

154. Smillie C, Garcillan-Barcia MP, Francia MV, Rocha EP, de la Cruz F. 2010. Mobility of plasmids. Microbiol Mol Biol Rev 74:434–52.

155. Hu B, Perepelov AV, Liu B, Shevelev SD, Guo D, Senchenkova SN, Shashkov AS, Feng L, Knirel YA, Wang L. 2010. Structural and genetic evidence for the close relationship between Escherichia coli O71 and Salmonella enterica O28 O-antigens. FEMS Immunol Med Microbiol 59:161–9.

156. Capps KM, Ludwig JB, Shridhar PB, Shi X, Roberts E, DebRoy C, Cernicchiaro N, Phebus RK, Bai J, Nagaraja TG. 2021. Identification, Shiga toxin subtypes and prevalence of minor serogroups of Shiga toxin-producing Escherichia coli in feedlot cattle feces. Sci Rep 11:8601.

157. King LA, Filliol-Toutain I, Mariani-Kurkidjian P, Vaillant V, Vernozy-Rozand C, Ganet S, Pihier N, Niaudet P, de Valk H. 2010. Family outbreak of Shiga toxin-producing Escherichia coli O123:H-, France, 2009. Emerg Infect Dis 16:1491–3.

158. Canchaya C, Fournous G, Brussow H. 2004. The impact of prophages on bacterial chromosomes. Molecular Microbiology 53:9–18.

159. Fortier LC, Sekulovic O. 2013. Importance of prophages to evolution and virulence of bacterial pathogens. Virulence 4:354–65.

160. Yano B, Taniguchi I, Gotoh Y, Hayashi T, Nakamura K. 2023. Dynamic changes in Shiga toxin (Stx) 1 transducing phage throughout the evolution of O26:H11 Stx-producing Escherichia coli. Sci Rep 13:4935.

161. Ramisetty BCM, Sudhakari PA. 2019. Bacterial ‘Grounded’ Prophages: Hotspots for Genetic Renovation and Innovation. Front Genet 10:65.

162. Jerse AE, Yu J, Tall BD, Kaper JB. 1990. A genetic locus of enteropathogenic Escherichia coli necessary for the production of attaching and effacing lesions on tissue culture cells. Proc Natl Acad Sci U S A 87:7839–43.

163. McDaniel TK, Jarvis KG, Donnenberg MS, Kaper JB. 1995. A genetic locus of enterocyte effacement conserved among diverse enterobacterial pathogens. Proc Natl Acad Sci U S A 92:1664–8.

164. Sperandio V, Kaper JB, Bortolini MR, Neves BC, Keller R, Trabulsi LR. 1998. Characterization of the locus of enterocyte effacement (LEE) in different enteropathogenic Escherichia coli (EPEC) and Shiga-toxin producing Escherichia coli (STEC) serotypes. FEMS Microbiol Lett 164:133–9.

165. Schmidt H, Hensel M. 2004. Pathogenicity islands in bacterial pathogenesis. Clin Microbiol Rev 17:14–56.

166. Stevens MP, Frankel GM. 2014. The Locus of Enterocyte Effacement and Associated Virulence Factors of Enterohemorrhagic Escherichia coli. Microbiol Spectr 2:Ehec-0007–2013.

167. Furniss RCD, Clements A. 2018. Regulation of the Locus of Enterocyte Effacement in Attaching and Effacing Pathogens. J Bacteriol 200.

168. Jores J, Rumer L, Wieler LH. 2004. Impact of the locus of enterocyte effacement pathogenicity island on the evolution of pathogenic Escherichia coli. International Journal of Medical Microbiology 294:103–113.

169. Sváb D, Falgenhauer L, Mag T, Chakraborty T, Tóth I. 2022. Genomic Diversity, Virulence Gene, and Prophage Arrays of Bovine and Human Shiga Toxigenic and Enteropathogenic Escherichia coli Strains Isolated in Hungary. Front Microbiol 13:896296.

170. Saile N, Schuh E, Semmler T, Eichhorn I, Wieler LH, Bauwens A, Schmidt H. 2018. Determination of virulence and fitness genes associated with the pheU, pheV and selC integration sites of LEE-negative food-borne Shiga toxin-producing Escherichia coli strains. Gut Pathog 10:43.

171. Dozois CM, Curtiss R, 3rd. 1999. Pathogenic diversity of Escherichia coli and the emergence of ‘exotic’ islands in the gene stream. Vet Res 30:157–79.

172. Lloyd AL, Rasko DA, Mobley HL. 2007. Defining genomic islands and uropathogen-specific genes in uropathogenic Escherichia coli. J Bacteriol 189:3532–46.

173. Memariani M, Najar-Peerayeh S, Zahraei Salehi T. 2019. Multi Locus VNTR (MLVA) Typing and Detection of the OI-122 Pathogenicity Island in Typical and Atypical Enteropathogenic Escherichia coli Isolated from Children with Acute Diarrhea. Arch Clin Infect Dis 14:e65855.

174. Ju W, Shen J, Toro M, Zhao S, Meng J. 2013. Distribution of pathogenicity islands OI-122, OI-43/48, and OI-57 and a high-pathogenicity island in Shiga toxin-producing Escherichia coli. Appl Environ Microbiol 79:3406–12.

175. Konczy P, Ziebell K, Mascarenhas M, Choi A, Michaud C, Kropinski AM, Whittam TS, Wickham M, Finlay B, Karmali MA. 2008. Genomic O island 122, locus for enterocyte effacement, and the evolution of virulent verocytotoxin-producing Escherichia coli. J Bacteriol 190:5832–40.

176. Desvaux M, Dalmasso G, Beyrouthy R, Barnich N, Delmas J, Bonnet R. 2020. Pathogenicity Factors of Genomic Islands in Intestinal and Extraintestinal Escherichia coli. Front Microbiol 11:2065.

177. Azpiroz MF, Bascuas T, Lavina M. 2011. Microcin H47 system: an Escherichia coli small genomic island with novel features. PLoS One 6:e26179.

178. Ziebell K, Johnson RP, Kropinski AM, Reid-Smith R, Ahmed R, Gannon VP, Gilmour M, Boerlin P. 2011. Gene cluster conferring streptomycin, sulfonamide, and tetracycline resistance in Escherichia coli O157:H7 phage types 23, 45, and 67. Appl Environ Microbiol 77:1900–3.

179. Partridge SR, Kwong SM, Firth N, Jensen SO. 2018. Mobile Genetic Elements Associated With Antimicrobial Resistance. Clinical Microbiology Reviews 31.

180. Haniford DB, Ellis MJ. 2015. Transposons Tn10 and Tn5. Microbiol Spectr 3:MDNA3-0002–2014.

181. Chopra I. 1978. Plasmid-determined tetracycline resistance in Escherichia coli K12: lack of evidence that resistance is related to changes in lipid metabolism [proceedings]. Biochem Soc Trans 6:431–3.

182. Aldema ML, McMurry LM, Walmsley AR, Levy SB. 1996. Purification of the Tn10-specified tetracycline efflux antiporter TetA in a native state as a polyhistidine fusion protein. Mol Microbiol 19:187–95.

183. Awosile B, Fritzler J, Levent G, Rahman MK, Ajulo S, Daniel I, Tasnim Y, Sarkar S. 2023. Genomic Characterization of Fecal Escherichia coli Isolates with Reduced Susceptibility to Beta-Lactam Antimicrobials from Wild Hogs and Coyotes. Pathogens 12.

184. Beck CM, Willett JL, Cunningham DA, Kim JJ, Low DA, Hayes CS. 2016. CdiA Effectors from Uropathogenic Escherichia coli Use Heterotrimeric Osmoporins as Receptors to Recognize Target Bacteria. PLoS Pathog 12:e1005925.

185. Bartelli NL, Passanisi VJ, Michalska K, Song K, Nhan DQ, Zhou H, Cuthbert BJ, Stols LM, Eschenfeldt WH, Wilson NG, Basra JS, Cortes R, Noorsher Z, Gabraiel Y, Poonen-Honig I, Seacord EC, Goulding CW, Low DA, Joachimiak A, Dahlquist FW, Hayes CS. 2022. Proteolytic processing induces a conformational switch required for antibacterial toxin delivery. Nat Commun 13:5078.

186. Heller DM, Tavag M, Hochschild A. 2017. CbtA toxin of Escherichia coli inhibits cell division and cell elongation via direct and independent interactions with FtsZ and MreB. PLoS Genet 13:e1007007.

187. Wang Q, Wei Y, Huang Y, Qin J, Liu B, Liu R, Chen X, Li D, Wang Q, Li X, Yang X, Li Y, Sun H. 2024. Z3495, a LysR-Type Transcriptional Regulator Encoded in O Island 97, Regulates Virulence Gene Expression in Enterohemorrhagic Escherichia coli O157:H7. Microorganisms 12.

188. Taschuk F, Cherry S. 2020. DEAD-Box Helicases: Sensors, Regulators, and Effectors for Antiviral Defense. Viruses 12.

189. Wang X, Kim Y, Ma Q, Hong SH, Pokusaeva K, Sturino JM, Wood TK. 2010. Cryptic prophages help bacteria cope with adverse environments. Nat Commun 1:147.

190. Camara B, Liu M, Reynolds J, Shadrin A, Liu B, Kwok K, Simpson P, Weinzierl R, Severinov K, Cota E, Matthews S, Wigneshweraraj SR. 2010. T7 phage protein Gp2 inhibits the Escherichia coli RNA polymerase by antagonizing stable DNA strand separation near the transcription start site. Proc Natl Acad Sci U S A 107:2247–52.

191. Li F, Cao L, Bahre H, Kim SK, Schroeder K, Jonas K, Koonce K, Mekonnen SA, Mohanty S, Bai F, Brauner A, Lee VT, Rohde M, Romling U. 2022. Patatin-like phospholipase CapV in Escherichia coli – morphological and physiological effects of one amino acid substitution. NPJ Biofilms Microbiomes 8:39.

192. Junkermeier EH, Hengge R. 2021. A Novel Locally c-di-GMP-Controlled Exopolysaccharide Synthase Required for Bacteriophage N4 Infection of Escherichia coli. mBio 12:e0324921.

193. Tsutakawa SE, Lafrance-Vanasse J, Tainer JA. 2014. The cutting edges in DNA repair, licensing, and fidelity: DNA and RNA repair nucleases sculpt DNA to measure twice, cut once. DNA Repair (Amst) 19:95–107.

194. Katsiou E, Nickel CM, Garcia AF, Tadros MH. 1999. Molecular analysis and identification of the radC gene from the phototrophic bacterium Rhodobacter capsulatus B10. Microbiol Res 154:233–9.

195. Liu B, Furevi A, Perepelov AV, Guo X, Cao H, Wang Q, Reeves PR, Knirel YA, Wang L, Widmalm G. 2020. Structure and genetics of Escherichia coli O antigens. FEMS Microbiol Rev 44:655–683.

196. Liu Y, Fratamico P, Debroy C, Bumbaugh AC, Allen JW. 2008. DNA sequencing and identification of serogroup-specific genes in the Escherichia coli O118 O antigen gene cluster and demonstration of antigenic diversity but only minor variation in DNA sequence of the O antigen clusters of E. coli O118 and O151. Foodborne Pathog Dis 5:449–57.

197. Liu B, Perepelov AV, Guo D, Shevelev SD, Senchenkova SN, Feng L, Shashkov AS, Wang L, Knirel YA. 2010. Structural and genetic relationships between the O-antigens of Escherichia coli O118 and O151. FEMS Immunol Med Microbiol 60:199–207.

198. Macnab R. 1996. Escherichia coli and Salmonella: cellular and molecular biology. Flagella and Motility:123–145.

199. Reid SD, Selander RK, Whittam TS. 1999. Sequence diversity of flagellin (fliC) alleles in pathogenic Escherichia coli. J Bacteriol 181:153–60.

200. Kalalah AA, Koenig SSK, Feng P, Bosilevac JM, Bono JL, Eppinger M. 2024. Pathogenomes of Shiga Toxin Positive and Negative Escherichia coli O157:H7 Strains TT12A and TT12B: Comprehensive Phylogenomic Analysis Using Closed Genomes. Microorganisms 12.

201. Ferdous M, Zhou K, Mellmann A, Morabito S, Croughs PD, de Boer RF, Kooistra-Smid AM, Rossen JW, Friedrich AW. 2015. Is Shiga Toxin-Negative Escherichia coli O157:H7 Enteropathogenic or Enterohemorrhagic Escherichia coli? Comprehensive Molecular Analysis Using Whole-Genome Sequencing. J Clin Microbiol 53:3530–8.

202. Wang L, Bai X, Ylinen E, Zhang J, Saxén H, Matussek A. 2023. Genetic Characterization of Intimin Gene (eae) in Clinical Shiga Toxin-Producing Escherichia coli Strains from Pediatric Patients in Finland. Toxins 15:669.

203. Elder JR, Fratamico PM, Liu Y, Needleman DS, Bagi L, Tebbs R, Allred A, Siddavatam P, Suren H, Gujjula KR, DebRoy C, Dudley EG, Yan X. 2020. A Targeted Sequencing Assay for Serotyping Escherichia coli Using AgriSeq Technology. Front Microbiol 11:627997.

204. Gelalcha BD, Brown SM, Crocker HE, Agga GE, Kerro Dego O. 2022. Regulation Mechanisms of Virulence Genes in Enterohemorrhagic Escherichia coli. Foodborne Pathog Dis 19:598–612.

205. Elliott SJ, Sperandio V, Girón JA, Shin S, Mellies JL, Wainwright L, Hutcheson SW, McDaniel TK, Kaper JB. 2000. The locus of enterocyte effacement (LEE)-encoded regulator controls expression of both LEE– and non-LEE-encoded virulence factors in enteropathogenic and enterohemorrhagic Escherichia coli. Infect Immun 68:6115–26.

206. Kolodziejek AM, Minnich SA, Hovde CJ. 2022. Escherichia coli 0157:H7 virulence factors and the ruminant reservoir. Curr Opin Infect Dis 35:205–214.

207. Oswald E, Schmidt H, Morabito S, Karch H, Marches O, Caprioli A. 2000. Typing of intimin genes in human and animal enterohemorrhagic and enteropathogenic Escherichia coli: characterization of a new intimin variant. Infect Immun 68:64–71.

208. Blanco M, Schumacher S, Tasara T, Zweifel C, Blanco JE, Dahbi G, Blanco J, Stephan R. 2005. Serotypes, intimin variants and other virulence factors of eae positive Escherichia coli strains isolated from healthy cattle in Switzerland. Identification of a new intimin variant gene (eae-eta2). BMC Microbiol 5:23.

209. Yang X, Sun H, Fan R, Fu S, Zhang J, Matussek A, Xiong Y, Bai X. 2020. Genetic diversity of the intimin gene (eae) in non-O157 Shiga toxin-producing Escherichia coli strains in China. Sci Rep 10:3275.

210. Cergole-Novella MC, Nishimura LS, Irino K, Vaz TM, de Castro AF, Leomil L, Guth BE. 2006. Stx genotypes and antimicrobial resistance profiles of Shiga toxin-producing Escherichia coli strains isolated from human infections, cattle and foods in Brazil. FEMS Microbiol Lett 259:234–9.

211. Franz E, Delaquis P, Morabito S, Beutin L, Gobius K, Rasko DA, Bono J, French N, Osek J, Lindstedt BA, Muniesa M, Manning S, LeJeune J, Callaway T, Beatson S, Eppinger M, Dallman T, Forbes KJ, Aarts H, Pearl DL, Gannon VP, Laing CR, Strachan NJ. 2014. Exploiting the explosion of information associated with whole genome sequencing to tackle Shiga toxin-producing Escherichia coli (STEC) in global food production systems. Int J Food Microbiol 187:57–72.

212. Robins-Browne RM, Holt KE, Ingle DJ, Hocking DM, Yang J, Tauschek M. 2016. Are Escherichia coli Pathotypes Still Relevant in the Era of Whole-Genome Sequencing? Front Cell Infect Microbiol 6:141.

213. Schutz K, Cowley LA, Shaaban S, Carroll A, McNamara E, Gally DL, Godbole G, Jenkins C, Dallman TJ. 2017. Evolutionary Context of Non-Sorbitol-Fermenting Shiga Toxin-Producing Escherichia coli O55:H7. Emerg Infect Dis 23:1966–1973.

214. Wang L, Rothemund D, Curd H, Reeves PR. 2003. Species-wide variation in the Escherichia coli flagellin (H-antigen) gene. J Bacteriol 185:2936–43.

215. Strachan NJ, Rotariu O, Lopes B, MacRae M, Fairley S, Laing C, Gannon V, Allison LJ, Hanson MF, Dallman T, Ashton P, Franz E, van Hoek AH, French NP, George T, Biggs PJ, Forbes KJ. 2015. Whole Genome Sequencing demonstrates that Geographic Variation of Escherichia coli O157 Genotypes Dominates Host Association. Sci Rep 5:14145.

216. Beutin L, Krause G, Zimmermann S, Kaulfuss S, Gleier K. 2004. Characterization of Shiga toxin-producing Escherichia coli strains isolated from human patients in Germany over a 3-year period. J Clin Microbiol 42:1099–108.

217. Bibbal D, Loukiadis E, Kerouredan M, Peytavin de Garam C, Ferre F, Cartier P, Gay E, Oswald E, Auvray F, Brugere H. 2014. Intimin gene (eae) subtype-based real-time PCR strategy for specific detection of Shiga toxin-producing Escherichia coli serotypes O157:H7, O26:H11, O103:H2, O111:H8, and O145:H28 in cattle feces. Appl Environ Microbiol 80:1177–84.

218. Gamage SD, Patton AK, Strasser JE, Chalk CL, Weiss AA. 2006. Commensal bacteria influence Escherichia coli O157:H7 persistence and Shiga toxin production in the mouse intestine. Infect Immun 74:1977–83.

219. Figler HM, Dudley EG. 2016. The interplay of Escherichia coli O157:H7 and commensal E. coli: the importance of strain-level identification. Expert Rev Gastroenterol Hepatol 10:415–7.

220. Baumler AJ, Sperandio V. 2016. Interactions between the microbiota and pathogenic bacteria in the gut. Nature 535:85–93.

221. Richter TKS, Michalski JM, Zanetti L, Tennant SM, Chen WH, Rasko DA. 2018. Responses of the Human Gut Escherichia coli Population to Pathogen and Antibiotic Disturbances. mSystems 3.

222. Wong CS, Jelacic S, Habeeb RL, Watkins SL, Tarr PI. 2000. The risk of the hemolytic-uremic syndrome after antibiotic treatment of Escherichia coli O157:H7 infections. N Engl J Med 342:1930–6.

223. Dundas S, Todd WT, Stewart AI, Murdoch PS, Chaudhuri AK, Hutchinson SJ. 2001. The central Scotland Escherichia coli O157:H7 outbreak: risk factors for the hemolytic uremic syndrome and death among hospitalized patients. Clin Infect Dis 33:923–31.

224. Gould LH, Demma L, Jones TF, Hurd S, Vugia DJ, Smith K, Shiferaw B, Segler S, Palmer A, Zansky S, Griffin PM. 2009. Hemolytic uremic syndrome and death in persons with Escherichia coli O157:H7 infection, foodborne diseases active surveillance network sites, 2000-2006. Clin Infect Dis 49:1480–5.

225. Foster DB. 2013. Modulation of the enterohemorrhagic E-coli virulence program through the human gastrointestinal tract. Virulence 4:315–323.

226. Johnson TJ, Nolan LK. 2009. Pathogenomics of the virulence plasmids of Escherichia coli. Microbiol Mol Biol Rev 73:750–74.

227. Messele YE, Trott DJ, Hasoon MF, Veltman T, McMeniman JP, Kidd SP, Djordjevic SP, Petrovski KR, Low WY. 2023. Phylogenetic Analysis of Escherichia coli Isolated from Australian Feedlot Cattle in Comparison to Pig Faecal and Poultry/Human Extraintestinal Isolates. Antibiotics (Basel) 12.

228. Wu R, Lv L, Wang C, Gao G, Yu K, Cai Z, Liu Y, Yang J, Liu JH. 2022. IS26-Mediated Formation of a Hybrid Plasmid Carrying mcr-1.1. Infect Drug Resist 15:7227–7234.

